# Quantification of hydrazine in biochemical assays and anammox bacteria using LC-MS

**DOI:** 10.64898/2026.02.09.704898

**Authors:** Femke J. Vermeir, Laura van Niftrik, Robert S. Jansen

**Author notes:** corresponding author: Robert S. Jansen.

## Abstract

Hydrazine is an industrially valuable product. Strikingly, hydrazine is also a key intermediate in the energy metabolism of anaerobic ammonium-oxidizing (anammox) bacteria, where it is formed by hydrazine synthase. To study the molecular mechanism and activity of isolated hydrazine synthase, a sensitive, relatively fast and easy method to quantify hydrazine is needed. However, reported methods such as colorimetric assays, MALDI-TOF MS, and enzymatic conversion of hydrazine to dinitrogen gas, are either insensitive or laborious. In this study, we describe the validation and application of a fast and simple liquid chromatography-mass spectrometry (LC-MS) method to reproducibly quantify hydrazine produced by anammox hydrazine synthase. Hydrazine was derivatized with benzaldehyde, and directly injected onto a C18 column coupled to a Q-TOF MS. To increase assay performance, ^15^N_2_-hydrazine was included as internal standard. The response ratio of hydrazine was linearly proportional to the hydrazine concentration from 0.05-1 µM with an average correlation coefficient of 0.9925. Intra- and inter-day accuracy lay between 88-113% and 95-105%, respectively. Intra- and inter-day precision (RSD, %) ≤ 11%. Hydrazine and derivatized hydrazine were stable when stored at -70°C or in the autosampler. We successfully applied the LC-MS method to determine hydrazine production by isolated hydrazine synthase and within cell lysate of anammox bacteria.

## Introduction

Hydrazine is an inorganic compound with the formula N_2_H_4_. It is a strong base, powerful reducing agent and is highly toxic [1-3]. Due to its properties, hydrazine is a commercially valuable product that is used, among others, to produce foaming agents, pesticides, pharmaceuticals, and monopropellant rocket fuel [4-8]. In nature, hydrazine is produced as a free intermediate in the energy metabolism of anaerobic ammonium-oxidizing (anammox) bacteria. These bacteria form dinitrogen gas from ammonium under anaerobic conditions with nitrite as final electron acceptor. The anammox metabolism comprises of three steps. First, nitrite is reduced to nitric oxide by nitrite reductase [9]. In the second step, the biochemically unique enzyme hydrazine synthase uses nitric oxide to activate ammonium and form hydrazine via the intermediate hydroxylamine [10-12]. Finally, hydrazine dehydrogenase oxidizes hydrazine to dinitrogen gas [13]. The electrons released and consumed in the anammox reaction are proposed to be transported in a cyclic manner via an electron transport chain and used to build a proton motive force across the membrane driving ATP synthesis [14, 15]. Although the production of the key intermediate hydrazine by hydrazine synthase is established, it is challenging to study the activity of the enzyme because hydrazine formation cannot be followed easily.

Previously, activity of isolated hydrazine synthase was researched either indirectly via the enzymatic production of dinitrogen gas, or directly via the quantification of hydrazine [12]. To measure hydrazine formation via dinitrogen gas, isolated hydrazine synthase was incubated with the substrates ammonium and nitric oxide in the presence of a second enzyme, hydroxylamine oxidoreductase (kustc1061), that converts hydrazine to dinitrogen gas. Although coupled enzyme reactions have been indispensable to study the formation of challenging enzyme products, several prerequisites must be met for correct results. Most importantly, the second reaction cannot be rate-limiting under any of the test conditions. Assuring this makes enzyme characterization under various conditions tedious. Moreover, hydroxylamine oxidoreductase also uses the anammox intermediate hydroxylamine as a substrate. Taken together, these drawbacks limit the use of this coupled assay for hydrazine synthase activity.

Direct measurements of hydrazine are complicated due to the reactive nature of the compound [16]. To overcome this challenge, hydrazine was previously derivatized with *para*-dimethylaminobenzaldehyde and detected with MALDI-TOF MS [12, 17]. Similarly, Oshiki et al. [18] published a comprehensive method to detect N-compounds such as hydrazine with MALDI-TOF MS to follow nitrogen transformation in bacterial cultures. However, these methods are relatively labor intensive, were not validated, and did not include an internal standard, limiting their robustness. Despite previous efforts, a sensitive, and relatively fast and easy method to quantify hydrazine production by hydrazine synthase is thus needed.

In recent years, various methods to detect low concentrations of hydrazine have been developed for other purposes. For instance, fluorescent probes were used to visualize and quantify hydrazine in cells [8, 19, 20]. However, a downside of this tool is that isotopes, which enable tracing atoms through the metabolic reaction, cannot be distinguished. Hydrazine has also frequently been measured with liquid chromatography-mass spectrometry (LC-MS). These methods rely on derivatization of hydrazine to a stable compound with a m/z that can be detected with MS. Most of the methods described are used to detect exposure to the suspected carcinogen hydrazine in humans.

Since hydrazine is mainly seen as a contaminant, the developed techniques focus on trace amounts of the compound. As a result, hydrazine has been measured in low concentrations starting from 94 pM (0.003 ng/mL) in, for example, pharmaceuticals [21], air [22, 23], drinking water and the human body [24-26]. In enzyme assays with isolated hydrazine synthase and bacterial cultures, hydrazine is produced in higher quantities (nM - µM range), shifting focus from low limits of detection to ease of use and robustness.

In this study, we validated a method for the measurement and quantification of hydrazine produced by the hydrazine synthase of anammox bacteria. This fast and simple method enables to reproducibly quantify hydrazine in small sample volumes relevant for enzyme assays, from 0.05 to 1 µM (1.6 - 32 ng/mL). Intra- and inter-day accuracy lay between 88-113% and between 95-105%, respectively (*n*=18). Intra- and inter-day precision (RSD, %) was less than 11% (*n*=18). Hydrazine and derivatized hydrazine can be stored at -70°C and in the autosampler of 8°C for at least 8 days. To demonstrate the suitability of our method, we successfully applied it to determine hydrazine production by isolated hydrazine synthase and in cell lysate of cultured anammox bacteria.

## Materials and Methods

### Reagents

Hydrazine sulfate salt (99% pure) was purchased from Sigma Aldrich and ^15^N_2_-hydrazine monosulfate (98% isotopic purity) was purchased from Cambridge Isotope Laboratories. For the derivatization of hydrazine, a derivatizing solution was prepared from benzaldehyde purchased from Sigma Aldrich Merck, acetonitrile hypergrade LC-MS purchased from Supelco Merck, and acetic acid purchased from Fisher chemicals. Formic acid was purchased from Fluka Honeywell and trifluoroacetic acid (99%) from Acros Organics. For the activity assays, hydroxylamine hydrochloride and ammonium chloride were purchased from Thermo Scientific.

### Preparation of stock and working solutions

For every analytical run, 1 mL hydrazine stock solution of 100 mM was freshly prepared in ultrapure water. For all samples, one internal standard stock solution of 1 mL 100 mM ^15^N_2_-hydrazine was prepared in ultrapure water, diluted to 0.1 µM and stored in aliquots at –70°C in between assays. One derivatizing stock solution was prepared consisting of 0.4% benzaldehyde (v/v) with 10 mM acetic acid in 70% acetonitrile and stored in a dark Greiner tube at room temperature in between assays. To test the effect of pH on derivatization, derivatizing solution was also prepared with 10 mM trifluoroacetic acid or 10 mM formic acid instead of 10 mM acetic acid.

### Preparation of calibration standards and validation samples

All hydrazine dilutions were made in ultrapure water. Calibration curves from 0.05 - 1 µM hydrazine were prepared fresh for every experiment. Quality controls comprised of a low concentration quality control of 0.3 µM hydrazine, a medium quality control of 0.5 µM hydrazine, and the high quality control of 0.8 µM hydrazine.

### Derivatization

Samples with hydrazine were always mixed in 1:1:2 (v/v/v) (internal standard:sample:derivatizing solution). Samples were immediately vortexed for 10 seconds and incubated for one hour in the dark at room temperature as was previously described by Cui et al [21]. Derivatizing solution diluted with ultrapure water (1:1 (v/v)) was used as blank in between measurements.

### LC-MS

Samples were subjected to LC-MS analysis performed by an Agilent 1290 II LC system with a 1290-series isocratic pump, multisampler, MCT, and high-speed pump coupled to an Agilent Accurate Mass 6546 Quadrupole Time of Flight (Q-TOF) instrument operated in the positive ionization mode. Sample vials with PTFE/silicone septa (Agilent) were used and the autosampler was set at 8°C. The isocratic pump continuously pumped a reference solution consisting of 1 µM Hexakis (1H, 1H, 3H-tetrafluoropropoxy) phosphazine, 5 µM purine, and 25 µM ammonium trifluoroacetate (Agilent technologies) at 2.5 mL/min, of which 1/100 was directed to the source via a splitter. The MS signals produced by reference compounds Hexakis and purine (m/z 121.0508 and 922.0997, respectively) were used for continuous mass calibration. For reversed phase chromatography, a Poroshell 120 EC-C18 column (2.1 × 50 mm, 1.9 μm, Agilent Infinity Lab) equipped with an Agilent 0.3 µm inline filter was used at 25°C and with a maximum pressure of 600 bar. A sample volume of 3 µL was injected onto the column, followed by separation with a 0.4 mL/min gradient of water (A) and acetonitrile (B) (both with 0.2% formic acid (Honeywell Fluka)): 30% B from 0-2 minutes, 30-100% B from 2-8 minutes, 100-30%B from 8-8.1 minutes, and finally re-equilibration of the column in 30% B from 8.1-10 minutes. The eluate was directed to waste for the first and final 2 minutes. After injection, the needle was washed with 70% methanol for 3 seconds (standard wash). Further settings used for the Dual AJS ESI were: gas temperature of 320°C, drying gas of 8 L/min, nebulizer gas 45 psi, sheath gas temperature of 350°C, flow of 11 L/min, Vcap was set to 3500 V, and nozzle voltage was 1000 V. The fragmentor was set at 125 V and the skimmer at 50 V. MS spectra were collected using full scan with a range of m/z 50-1200 at 1000 ms/spectrum. Collected data was analyzed using Quant software 10.0 from Agilent (Agilent Technologies). Signals for hydrazine and ^15^N_2_-hydrazine (m/z 209.1074 and m/z 211.1014, respectively), were analyzed with a 10-ppm window and peak smoothing (function width 15, Gaussian width 5).

### Assay validation

#### Linearity

To evaluate linearity, 6 calibration curves consisting of 12 concentration levels (0.05, 0.08, 0.1, 0.15, 0.2, 0.3, 0.4, 0.5, 0.7, 0.8, 0.9, and 1 µM hydrazine) were measured. Each concentration was prepared and assayed in duplicate on three separate days. Calibration curves were obtained by plotting the peak area of hydrazine divided by the peak area of ^15^N_2_-hydrazine versus concentration. With Quant software (Agilent technologies), the linearity of the calibration curve was determined with a weighting of 1/x^2^. Accuracy was calculated by dividing the calculated concentration of hydrazine in the sample by the expected concentration. Datapoints for the LLOQ were excluded when the accuracy deviated > 20%. The other calibration levels were excluded when the deviation was > 15%.

#### Accuracy and precision

Accuracy was calculated by dividing the calculated hydrazine concentration in a spiked sample by the nominal hydrazine concentration for that sample. Intra-day accuracy was defined as the mean of the accuracy-values per validation run. Inter-day accuracy was defined as the mean of all accuracy values. Precision was calculated as the coefficient of variation (RSD, %), in which the standard deviation is divided by the mean hydrazine concentration. For intra-day precision the mean and standard deviation of the hydrazine concentration per validation run was used. For inter-day precision the mean and standard deviation of the hydrazine concentration in all samples was used.

#### Carry-over

Carry-over of hydrazine was assessed for the blank following the standard level ULQ. Carry-over was calculated by dividing the area hydrazine in the blank by the area hydrazine in LLOQ sample. As a cut-off, 20% of the area of hydrazine in the LLOQ level was used. For ^15^N_2_-hydrazine, the area of ^15^N_2_-hydrazine in the blank was divided by the area of ^15^N_2_-hydrazine in the LLOQ. For this area, 5% of the area in calibration levels was set as cut-off.

#### Stability of hydrazine

To test the stability of hydrazine solution in different conditions, samples of 0.3 µM and 0.8 µM hydrazine were stored at -70°C, -20°C, 4°C, in the autosampler of the LC-MS system (8°C), or at room temperature (∼ 20°C). The different samples were mixed with derivatizing solution (1:2 (v/v)) and 0.1 µM ^15^N_2_-hydrazine was added (1:4 (v/v)) before measuring hydrazine concentrations with LC-MS. To test the stability of derivatized hydrazine, samples with 0.3 µM and 0.8 µM hydrazine were mixed 1:2 (v/v) with derivatizing solution prior to storage at - 70°C, -20°C, 4°C, in the autosampler of the LC-MS system (8°C), or at room temperature (∼ 20°C). Samples were measured twice: at 8 days of storage and at 15 days of storage. Before measuring the samples on the LC-MS, 0.1 µM ^15^N_2_-hydrazine was added in 1:4 (v/v). For all samples accuracy and precision were calculated.

#### Cultivation of anammox bacteria

The anammox model species ‘*Candidatus* Kuenenia stuttgartiensis’ MBR1 was cultivated in an enrichment culture in a 12 liter single cell membrane bioreactor as previously described by Kartal et al. [27]. In brief, the reactor was operated at 33°C and kept anoxic by continuous flushing of the reactor itself and medium vessel with Ar/CO_2_ (95/5%, 10 mL min^−1^). CO_2_ in the supplied gas was sufficient to maintain the pH in the reactors between 7.0 and 7.4. Excess biomass was removed at 1.1 L per day, resulting in a doubling time of 10 days. Medium consisted of: 45 mM (NH_4_)_2_SO_4_, 45 mM NaNO_2_, 10 mM KHCO_3_, 0.2 mM NaH_2_PO_4_, 0.6 mM HCl, 1 mM CaCl_2_·2H_2_O, 0.4 mM MgSO_4_·7H_2_O and 22.5 µM FeSO_4_·7 H_2_O, and trace elements 0.625 µM CoCl_2_·6H_2_O, 0.625 µM CuSO_4_·5H_2_O, 0.125 µM H_3_BO_3,_ 3.125 µM MnCl_2_·4H_2_O, 0.563 µM Na_2_MoO_4_·2H_2_O, 0.125 µM Na_2_WO_4_·2H_2_O, 0.5 µM NiCl_2_·6H_2_O, 0.375 µM SeO_2_, and 0.938 nM ZnSO_4_·7H_2_O.

#### Isolation of hydrazine synthase

Hydrazine synthase was isolated from anammox bacteria via a method adapted from Kartal et al. [27]. All transfer and purification steps were carried out under anaerobic conditions (glove box atmosphere of 100% dinitrogen gas) at 16°C. In brief, 700 ml collected bioreactor culture was centrifuged and the pellet was resuspended in 20 mM Tris pH 7.3. Then, cells were broken by passing them once through a French press cell at 138 MPa (American Instrument Company). Membranes and whole cells were pelleted by centrifugation and soluble proteins were loaded onto a 70 ml Q-Sepharose column (XK 26/20, GE Healthcare/Cytiva) equilibrated with 20 mM Tris pH 7.3, connected to an Äkta purifier (GE Healthcare/Cytiva). Hydrazine synthase eluted isocratically with 200 mM NaCl. Proteins were buffer exchanged to 20 mM potassium phosphate buffer pH 7.0 and concentrated on a 50 kDa cutoff Amicon pressure filter unit. The sample was loaded onto a hydroxy apatite column (CHTII, 40 µm particle size, GE Healthcare/Cytiva, 15 mL, packed in house) equilibrated with 20 mM potassium phosphate buffer pH 7.0. Hydrazine synthase eluted at 150 mM potassium phosphate and was concentrated on 30 kDa spinfilters (Sartorius).

Purification of hydrazine synthase was verified with sodium dodecyl sulfate polyacrylamide gel electrophoresis (SDS-PAGE) and changes in absorption typical to hydrazine synthase were followed with UV-visible spectroscopy (Cary 60 spectrophotometer, Agilent) as previously described by Dietl et al. [11]. Hydrazine synthase concentration in µg/µL was calculated from the absorbance at wavelength 280 nm measured with a Cary 60 spectrophotometer (Agilent) and calculated with the formula: concentration (µg/µL) = (A_280_ / ε_280_ of hydrazine synthase) × molecular weight of hydrazine synthase. In which the 280 nm extinction coefficient is 26,7250 M^- 1^cm^-1^, and the molecular weight of hydrazine synthase is 160,997.73 Da. Hydrazine synthase was stored anaerobically at -20°C until used.

#### Application of the LC-MS method to measure activity of isolated hydrazine synthase

A final amount of 15 µg HZS was incubated with 1 mM ammonium chloride (99%, Thermo Fisher) and 10 µM hydroxylamine hydrochloride (99%, Thermo Scientific) in an end-volume of 500 µL 20 mM potassium phosphate buffer pH 7.0. Hydrazine production was measured over time via the addition of 25 µL reaction mixture and 25 µL internal standard (^15^N_2_-hydrazine) to 50 µL derivatizing solution every 30 seconds for 2 minutes, at 3.5 minutes and once more at 5 minutes. Samples were incubated in the dark for at least one hour for the derivatization of hydrazine with benzaldehyde to form 1,2-dibenzylidenehydrazine. All assays were performed in anaerobic conditions. Samples were centrifuged at 20817 x *g* for 5 minutes at 15°C to pellet denatured HZS prior to transferring the sample to a vial used for LC-MS analysis.

Control samples were measured for every assay. To this end, 1 µL of HZS stock solution, 99 µL potassium phosphate buffer supplemented with hydroxylamine and ammonium, and 100 µL internal standard were added to 200 µL derivatizing solution.

#### Application of the LC-MS method to measure hydrazine in anammox bacteria lysate

Samples of 1 mL were taken from the *K. stuttgartiensis* bioreactor and immediately placed on ice. To obtain a cell pellet, the bacteria were centrifuged at 20817 x *g* for 5 minutes at 4°C. The cell pellet was resuspended in derivatizing solution (1:3 (v/v)) and internal standard was added (1:4 (v/v)). Samples were measured with LC-MS after a one-hour incubation in the dark at room temperature.

#### Figures

Figures were made in Rstudio version 4.4.1 with the ggpubr, ggpmisc, ggpp, ggplot2, cowplot, and dplyr packages, and in Chemdraw 22.0.0.

## Results & Discussion

### Derivatization reaction and optimization

Hydrazine is a low molecular weight compound (32.0452 g/mol) that is challenging to detect with LC-MS. Therefore, we derivatized hydrazine with benzaldehyde to the stable product 1,2-dibenzylidenehydrazine (208.26 g/mol) (Fig. 1). Benzaldehyde reacts easily with hydrazine, and 1,2-dibenzylidenehydrazine is readily ionized by electron spray ionization (ESI) in positive ionization mode [21]. The reaction between hydrazine and benzaldehyde is often carried out in acidic conditions to ensure that benzaldehyde reacts with the amino groups of hydrazine, and to catalyze the loss of hydrogen to drive the reaction towards the formation of the derivatized product. Cui et al. previously tested different concentrations of benzene sulfonic acid, benzoic acid and formic acid, and concluded that the reaction was most rapid in a benzoic acid buffer at pH 3.74 [21]. Here, we assessed hydrazine derivatization in three different LC-MS-compatible derivatizing solutions: acetic acid pH 5, formic acid pH 4, and trifluoroacetic acid pH 2. After one hour of incubation, there were only minor differences between the product signal in these three solutions (Table 1 and Table S1). These results indicate that the pH between 2-5 is not a critical parameter as long as acidity is maintained. Since acetic acid is the mildest acid tested, we added it to buffer the derivatizing solution and ensure that the proper acidity is sustained.

**Table 1.** Hydrazine derivatization results in various acids. The addition of three different acids to derivatizing solution, 10 mM acetic acid (AA), formic acid (FA) or trifluoroacetic acid (TFA), were tested. Data are presented as mean ± SD (*n*=3).

| Derivatizing solution | pH | Response 0.3 μM hydrazine | Response 0.8 μM hydrazine |
| --- | --- | --- | --- |
| | | mean (counts) ± SD | mean (counts) ± SD |
| derivatizing solution | 5 | 2.3E+05 ± 8.3E+03 | 6.2E+05 ± 7.2E+03 |
| derivatizing solution + 10 mM AA | 5 | 2.0E+05 ± 5.7E+03 | 5.7E+05 ± 1.1E+04 |
| derivatizing solution + 10 mM FA | 4 | 2.0E+05 ± 6.7E+03 | 5.9E+05 ± 2.4E+04 |
| derivatizing solution + 10 mM TFA | 2 | 2.0E+05 ± 1.5E+04 | 5.8E+05 ± 1.2E+04 |

**Fig. 1.**
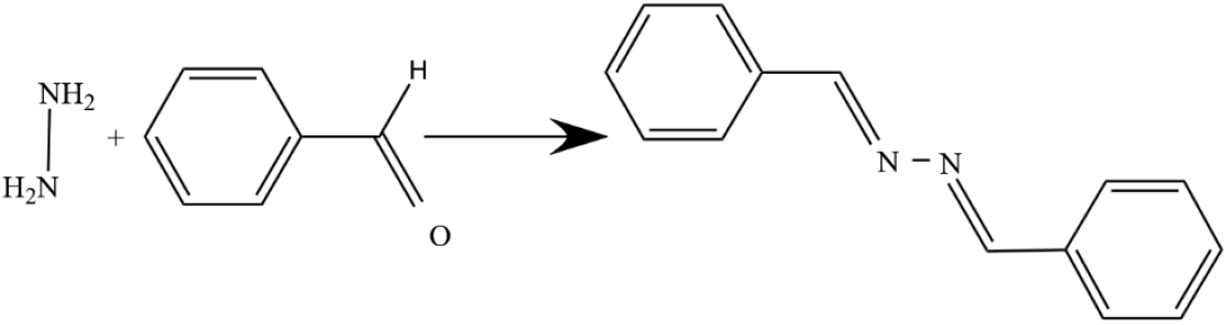
Reaction between hydrazine and benzaldehyde to 1,2-dibenzylidenehydrazine. Hydrazine can be measured with LC-MS after derivatization.

### Chromatography and internal standard

To separate the derivatized hydrazine from potential interferences, we subjected it to C18 chromatography and found that a simple gradient of acetonitrile in water (both with 0.2% formic acid) resulted in sharp, well-shaped peaks (Fig. 2). For extra robustness of the method, ^15^N_2_-hydrazine was added as internal standard. As expected, the natural and stable isotope-labeled hydrazine co-eluted (Fig. 2). Although the M+2 peak of unlabeled derivatized hydrazine produces a detectable signal at m/z 211, the accurate mass of this signal deviates 52 ppm from the m/z of derivatized ^15^N_2_-hydrazine. At the applied 10 ppm extraction window, the natural isotope signal of derivatized ^14^N_2_-hydrazine was therefore not detected in the derivatized ^15^N_2_-hydrazine mass extraction window (Fig. 2).

**Fig. 2.**
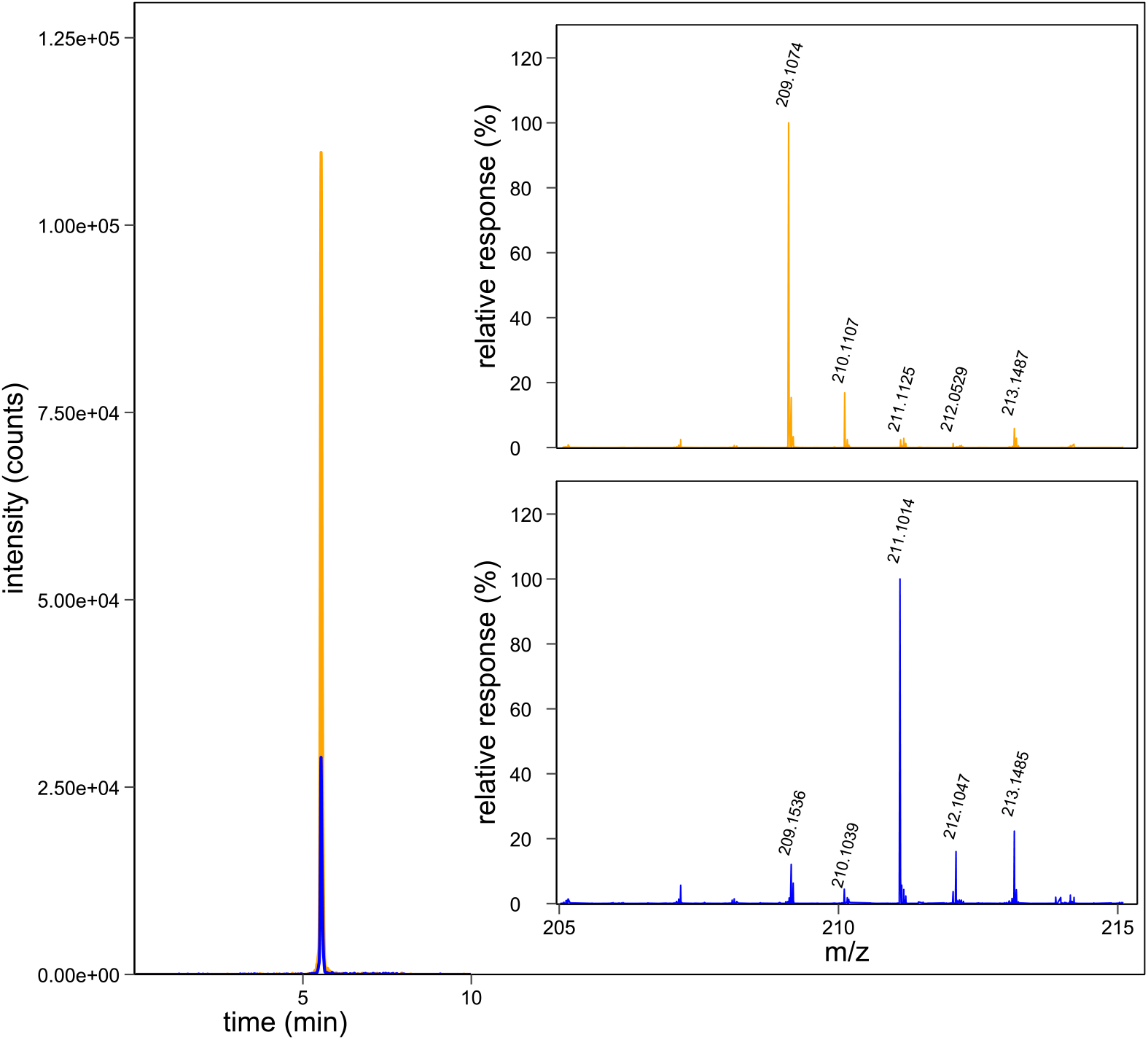
Derivatized hydrazine was separated from potential interferences via C18 chromatography. The representative peak for hydrazine (0.3 µM) in orange and ^15^N_2_-hydrazine (0.1 µM) in blue shows that the natural and stable isotope-labeled hydrazine co-eluted as sharp, well-shaped peaks. The MS spectrum of the hydrazine signal in the ULQ without internal standard (upper inset, in orange) shows that the ^15^N_2_-hydrazine signal is absent (signal at m/z 211.1014). The MS spectrum of the internal standard shows that the unlabeled hydrazine signal (m/z 209.1074) is similarly absent in the zerocalibrator (bottom inset, in blue). Relative response represents the intensity of the compound signal compared to the intensity of either unlabeled hydrazine (m/z 209.1074) or ^15^N_2_-hydrazine (m/z 211.1014). (*n*=1).

### Method validation

#### Linearity and sensitivity

To assess the linearity and sensitivity of the method, calibration samples were injected to construct two calibration curves per analytical run, on three separate days. The calibration curves consisted of 12 peak area ratios of derivatized hydrazine to derivatized ^15^N_2_-hydrazine and were fitted by linear regression with 1/x^2^ weighting. The calibration curves were linear over a range of 0.05-1 µM hydrazine, with an average coefficient of determination (R^2^) of 0.9925 and an average line equation of 11.21636 * x + 0.107197 (*n*=6) (Fig. 3 and Table S1). The hydrazine signal in the LLOQ showed a high signal to noise ratio (Fig. 4).

**Fig. 3.**
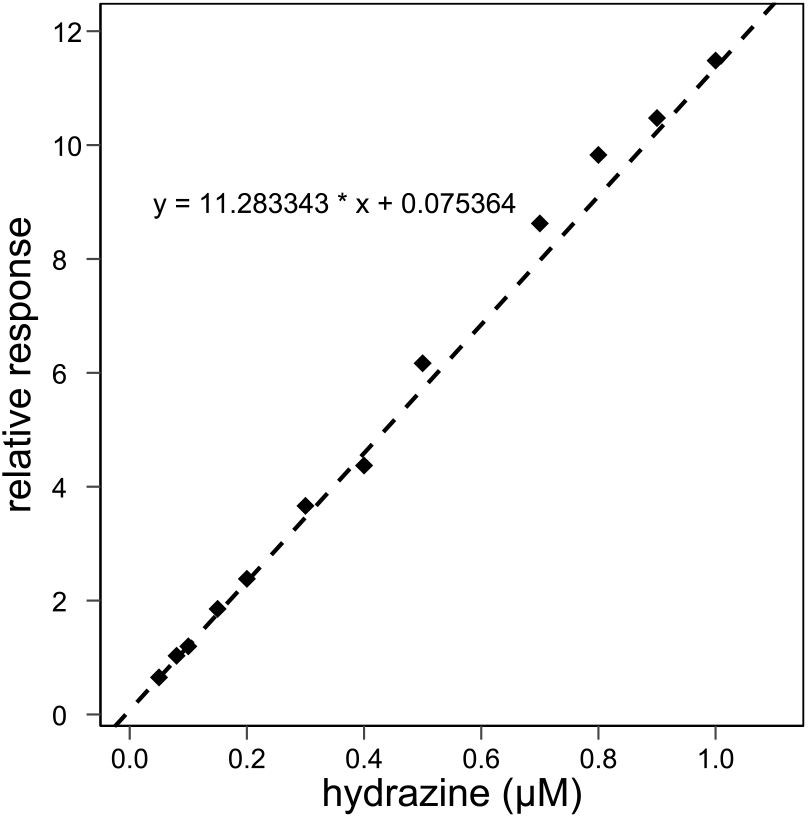
Representative calibration curve. Calibration curves consisted of 12 calibrator levels from 0.05-1 µM hydrazine supplemented with 0.1 µM ^15^N_2_-hydrazine as internal standard. Relative response (the peak area ratio of derivatized hydrazine to derivatized ^15^N_2_-hydrazine) was fitted by linear regression with 1/x^2^ weighting (*n*=1).

**Fig. 4.**
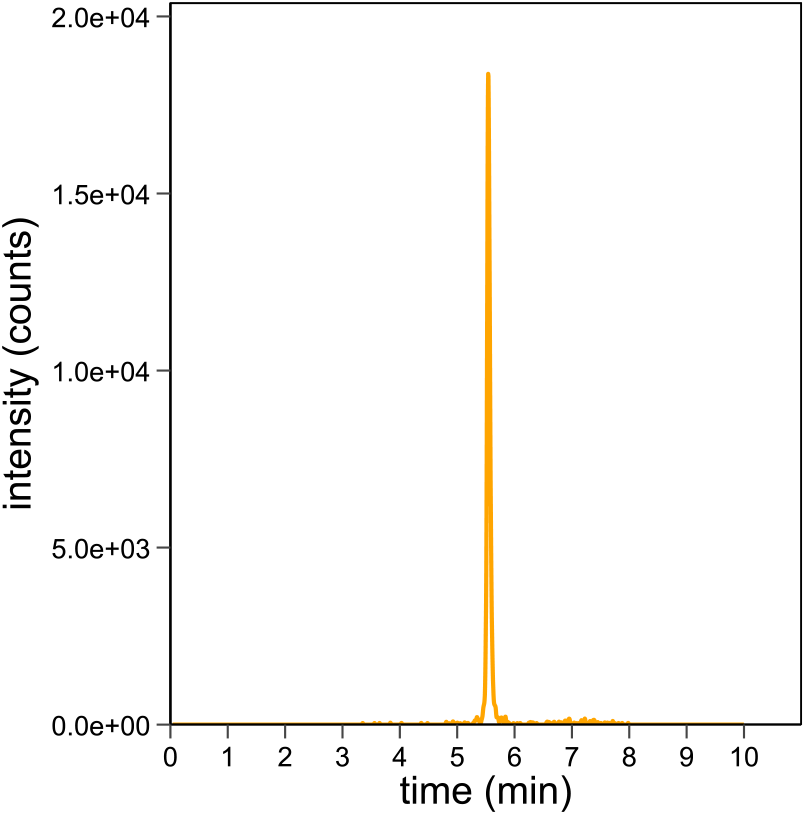
Representative chromatogram of hydrazine in an LLOQ sample containing 0.05 µM hydrazine (*n=1*).

#### Carry-over

During validation, carry-over was assessed in the blank following the calibration standard at the ULQ [28]. Carry-over of hydrazine was calculated as the peak area hydrazine in the blank divided by the peak area hydrazine in the LLOQ and should not exceed 20%. For the internal standard, the peak area for ^15^N_2_-hydrazine in the blank should be lower than 5% of the ^15^N_2_-hydrazine peak area in the LLOQ. The carry-over determined in analytical run 1 was below 8% for hydrazine and below 3% for ^15^N_2_-hydrazine (Table S1). Thus, the carry-over was within limits. Even though the influence of the carry-over was acceptable when assessed, we noticed carry-over in other analytical runs and thus advise to include blanks for monitoring.

#### Accuracy and precision

To further validate the method, four quality controls of LLOQ, low, medium, and high hydrazine concentrations were analyzed from six replicates on three different days, based on ICH guideline M10 on bioanalytical method validation [28] (Table 2). Of the quality controls (*n*=18), 93% fell within 85-115% accuracy and per concentration level ≥ 89% of the quality controls fell within these limits. Accuracy (relative error) was between 88 and 113% and between 95 and 105% intra- and inter-day, respectively (*n*=18). Intra- and inter-day precision (RSD, %) lay between 1.5 and 6.9% and between 7.1 and 11% respectively (*n*=18) (Table S1).

**Table 2.** Assay performance. The hydrazine quantification method was validated with four quality controls of LLOQ, low, medium, and high hydrazine concentrations based on the ICH guideline on bioanalytical method validation. Data are presented as mean ± SD (*n*=6).

| Quality control<br>($\mu\text{M}$) | Mean ± SD<br>($\mu\text{M}$) | Intra-day | | | | | | Inter-day | |
| --- | --- | --- | --- | --- | --- | --- | --- | --- | --- |
|  |  | Accuracy<br>(%) |  |  | Precision<br>(%) |  |  | Accuracy<br>(%) | Precision<br>(%) |
|  |  | Run 1 | Run 2 | Run 3 | Run 1 | Run 2 | Run 3 |  |  |
| 0.05 | 0.049 ± 0.004 | 89.9 | 101.9 | 101.6 | 3.6 | 4.4 | 6.2 | 97.8 | 7.6 |
| 0.3 | 0.313 ± 0.003 | 90.8 | 109.7 | 113.0 | 6.0 | 1.5 | 6.9 | 104.5 | 10.8 |
| 0.5 | 0.493 ± 0.035 | 91.0 | 101.9 | 103.4 | 4.6 | 1.9 | 5.1 | 98.6 | 7.1 |
| 0.8 | 0.762 ± 0.056 | 88.0 | 97.3 | 100.6 | 3.4 | 6.7 | 3.1 | 95.3 | 7.3 |

#### Stability

To assess the stability of hydrazine and derivatized hydrazine, their stability was evaluated under various common laboratory storage conditions at 8 and 15 days. Triplicates of 0.3 µM or 0.8 µM hydrazine were either dissolved in ultrapure water or derivatized with benzaldehyde 1:2 (v/v), and stored at -70°C, -20°C, 4°C, in the autosampler of the LC-MS system (8°C) or at room temperature. Samples were stored in the dark because benzaldehyde is light sensitive. Analytes were considered stable when the accuracy fell within 85-115% of the nominal value. Hydrazine solutions in ultrapure water were found to be stable in all conditions for 8 days, except for samples with 0.3 µM hydrazine stored at -20°C or at room temperature. Hydrazine solution can be stored at -70°C, 4°C, and the autosampler of 8°C up to 15 days (Fig. 5a and 5b). The processed samples with 0.3 µM derivatized hydrazine could be stored for 15 days at -70°C and in the autosampler. Processed samples with 0.8 µM derivatized hydrazine were stable for 8 days at -70°C or in the autosampler (Fig. 5c and 5d) (Table S1). The unexpected instability of samples with derivatized hydrazine stored at -20°C is possibly due to phase separation in the 70% acetonitrile solution. Acetonitrile freezes at -45°C, thus in samples stored at -20 °C frozen water can separate from the still soluble acetonitrile which could affect the chemical properties of the solution [29, 30].

**Fig. 5.**
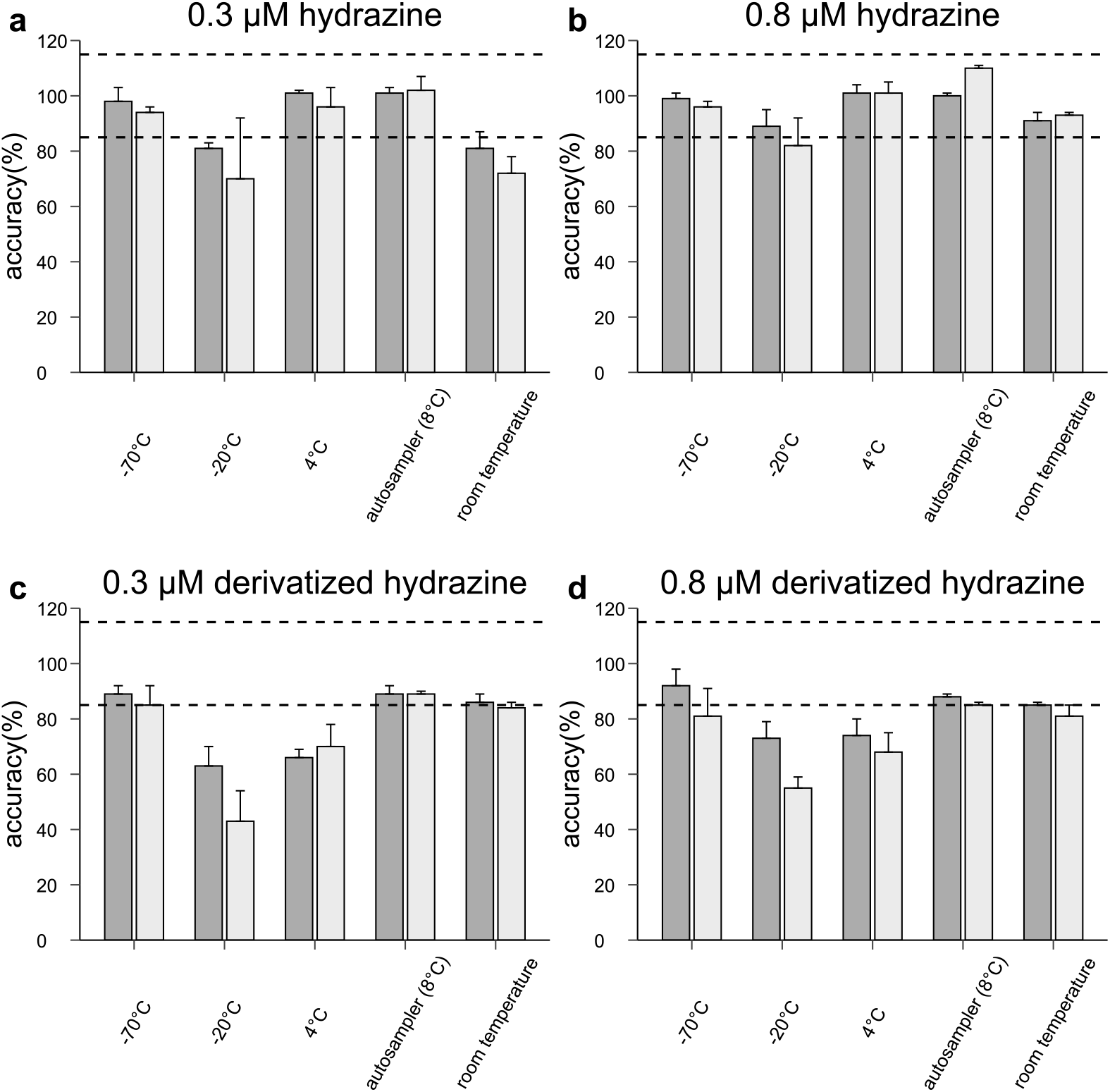
Sample stability. Stability of hydrazine (a and b) and derivatized hydrazine in solution (c and d) at 8 and 15 days of storage, as measured with the validated LC-MS method. Data are presented as mean ± SD (*n*=3), accuracy limits for stability of the samples is represented with a dotted line: for the lower limit 85% and upper limit 115%, data measured on day 8 is represented in dark grey and data measured on day 15 is represented in light grey.

#### Application

To demonstrate the applicability of the LC-MS method, it was applied to an *in vitro* enzyme assay with HZS and to anammox bacteria lysate. For the enzyme assay, HZS was isolated from *K. stuttgartiensis* and incubated with substrates ammonium and hydroxylamine. The hydrazine formation was followed with our method over the course of 6 minutes (Fig. 6 and Table S1). Our method revealed hydrazine formation with levels up to 1.01 ± 0.05 µM. For the application of the method to anammox bacteria, the bacteria were collected from a bioreactor culture and lysed. Our assay showed that this bacterial lysate contained 0.20 ± 0.02 µM hydrazine (Fig. 7 and Table S1). Thus, this method enables the detection of hydrazine production in enzymatic assays with HZS or within anammox bacteria lysate. The internal standard of ^15^N_2_-hydrazine allows for reliable quantification of hydrazine production in complex biological samples. Moreover, isotopic labelling studies are also possible when ^15^N_2_-hydrazine is not used as an internal standard. This leads to new possibilities in the research on the molecular mechanism and activity of HZS and its role in the production of the unusual intermediate hydrazine in the anammox metabolism.

**Fig. 6.**
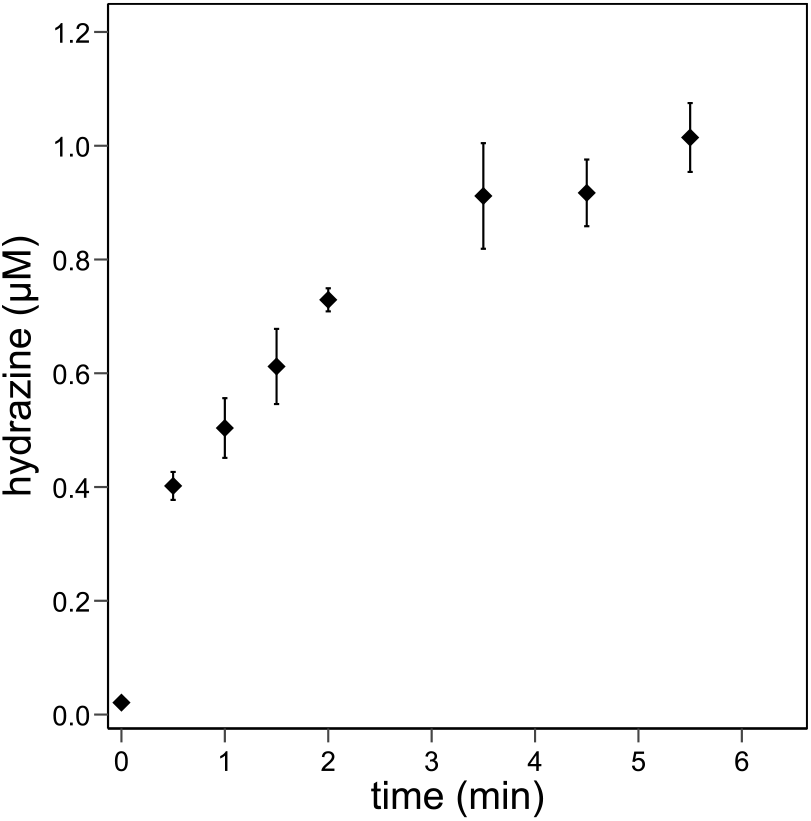
Hydrazine production by HZS isolated from “*Candidatus* Kuenenia stuttgartiensis” bacteria as measured with the validated LC-MS method. Activity assays contained 15 µg HZS and 1 mM ammonium and 10 µM hydroxylamine in potassium phosphate buffer. Data represented as mean ± SD (*n*=3).

**Fig. 7.**
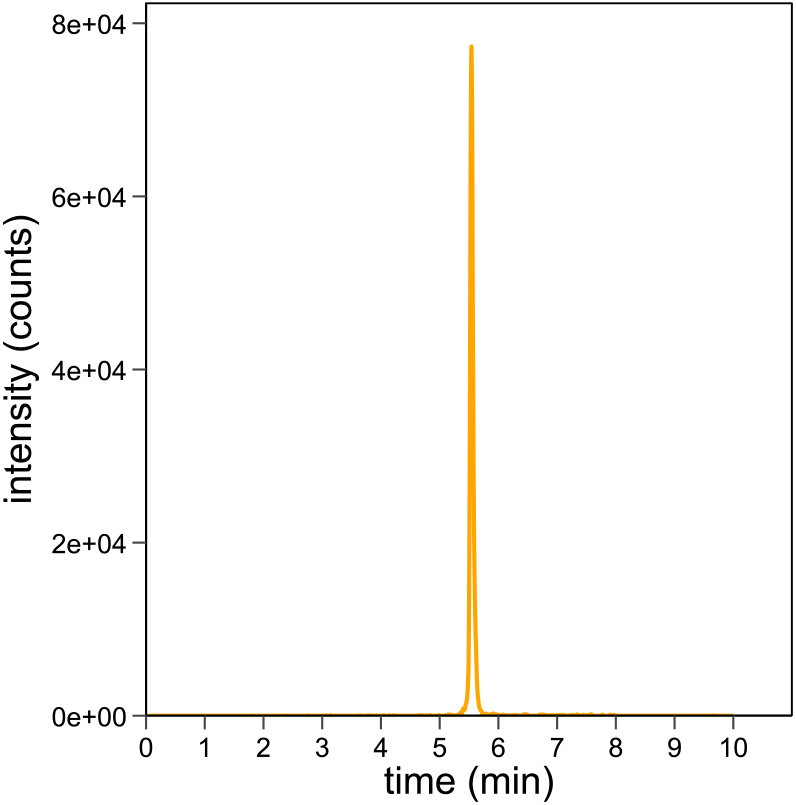
Representative chromatogram of hydrazine in lysate of anammox bacterium “*Candidatus* Kuenenia stuttgartiensis” as measured with the validated LC-MS method. The concentration was determined to be 0.20 ± 0.02 µM (*n*=3).

## Conclusion

In conclusion, we validated a method offering a practical and reliable approach for measuring hydrazine production, balancing robustness and ease of use with acceptable sensitivity and accuracy. The method enables to reproducibly quantify hydrazine from 0.05 to 1 µM (1.6 - 32 ng/mL) in small sample volumes relevant for enzyme assays. Furthermore, hydrazine and derivatized hydrazine are stable when stored at -70°C or in the autosampler of 8°C for at least 8 days. Finally, we demonstrated that the hydrazine quantification method is suitable for determination of hydrazine production by isolated hydrazine synthase and in cell lysate of cultured anammox bacteria.

## Declarations

The authors declare that they do not have financial or non-financial interests that are directly or indirectly related to the work submitted for publication.

## Acknowledgements

We would like to thank Guylaine Nuijten for assistance and co-maintenance of anammox bioreactor systems and Wouter Versantvoort, Rob de Graaf, and Jona Merx for helpful discussions.

This work was supported by the Netherlands Organisation for Scientific Research (NWO) [VI.Vidi.192.001] awarded to LvN.

## Data availability

The data of this study are available on request.

## Author contributions

FJV performed the experiments and analyzed the data, RSJ and LVN supervised the work. All authors contributed to writing of the manuscript.

## Ethics, Consent to Participate, and Consent to Publish declarations

Not applicable

