## Supplementary material for "Quantification of hydrazine in biochemical assays and anammox bacteria using LC-MS": Table S1

Quantification of hydrazine in biochemical assays and anammox bacteria using LC-MS
Analytical Sciences
Femke J. Vermeir[a], Laura van Niftrik[a], Robert S. Jansen[a*]
[a] Department of Microbiology, Radboud Institute for Biological and Environmental Sciences, Faculty of Science, Radboud University, Heyendaalseweg 135, 6525 AJ Nijmegen, The Netherlands

**Supplementary table 1** - Data obtained with LC-MS analyzed with Quant software 10.0 (Agilent Technologies) with a 10-ppm window and peak smoothing (function width 15, Gaussian width 5). Acq. Date-Time = acquired date and time, Exp. Conc. = expected concentration in µM, Resp. = response, Calc. Conc. = calculated concentration in µM, Final Conc. = final concentration in µM, RT = retention time in minutes.

| | | | | hydrazine Method | hydrazine Results | | | | | $^{15}N_2$-hydrazine (ISTD) Results | |
|---|---|---|---|---|---|---|---|---|---|---|---|
| **Name** | **Type** | **Level** | **Acq. Date-Time** | **Exp. Conc.** | **RT** | **Resp.** | **Calc. Conc.** | **Final Conc.** | **Accuracy** | **RT** | **Resp.** |
| **validation run 1** | | | | | | | | | | | |
| blank1 | DoubleBlank | | 10/30/2023 18:02 | | | | | | | 5.528 | 632 |
| blank2 | DoubleBlank | | 10/30/2023 18:13 | | 5.542 | 914 | 0.1028 | 0.1028 | | 5.504 | 740 |
| blank3 | DoubleBlank | | 10/30/2023 18:24 | | 5.533 | 801 | | | | | |
| zerocalibrator1 | Sample | | 10/30/2023 18:34 | | 5.539 | 521 | 0 | 0 | | 5.539 | 123914 |
| L1_1 | Cal | 0.05 | 10/30/2023 18:45 | 0.05 | 5.54 | 73528 | 0.051 | 0.051 | 101.9 | 5.54 | 113048 |
| L2_1 | Cal | 0.08 | 10/30/2023 18:56 | 0.08 | 5.539 | 114528 | 0.0847 | 0.0847 | 105.8 | 5.539 | 111123 |
| L3_1 | Cal | 0.1 | 10/30/2023 19:06 | 0.1 | 5.545 | 113772 | 0.0993 | 0.0993 | 99.3 | 5.541 | 95099 |
| L4_1 | Cal | 0.15 | 10/30/2023 19:17 | 0.15 | 5.54 | 227870 | 0.1576 | 0.1576 | 105.1 | 5.54 | 122910 |
| L5_1 | Cal | 0.2 | 10/30/2023 19:27 | 0.2 | 5.543 | 264344 | 0.2045 | 0.2045 | 102.2 | 5.539 | 110958 |
| L6_1 | Cal | 0.3 | 10/30/2023 19:38 | 0.3 | 5.54 | 408444 | 0.318 | 0.318 | 106 | 5.54 | 111488 |
| L7_1 | Cal | 0.4 | 10/30/2023 19:49 | 0.4 | 5.54 | 496781 | 0.3808 | 0.3808 | 95.2 | 5.54 | 113613 |
| L8_1 | Cal | 0.5 | 10/30/2023 19:59 | 0.5 | 5.541 | 711545 | 0.54 | 0.54 | 108 | 5.541 | 115362 |
| L9_1 | Cal | 0.7 | 10/30/2023 20:10 | 0.7 | 5.543 | 919441 | 0.7576 | 0.7576 | 108.2 | 5.539 | 106614 |
| L10_1 | Cal | 0.8 | 10/30/2023 20:21 | 0.8 | 5.541 | 1061491 | 0.8643 | 0.8643 | 108 | 5.541 | 108017 |
| L11_1 | Cal | 0.9 | 10/30/2023 20:31 | 0.9 | 5.538 | 1191554 | 0.9217 | 0.9217 | 102.4 | 5.538 | 113747 |
| L12_1 | Cal | 1 | 10/30/2023 20:42 | 1 | 5.541 | 1368351 | 1.0111 | 1.0111 | 101.1 | 5.541 | 119148 |
| preblank6 | DoubleBlank | | 10/30/2023 20:53 | | 5.561 | 5788 | 0.1325 | 0.1325 | | 5.523 | 3686 |
| preblank7 | DoubleBlank | | 10/30/2023 21:03 | | 5.534 | 2468 | 0.2775 | 0.2775 | | 5.534 | 770 |
| LLOQ_1 | QC | 0.05 | 10/30/2023 21:14 | 0.05 | 5.539 | 68295 | 0.0446 | 0.0446 | 89.3 | 5.543 | 117939 |
| Low_QC_1 | QC | 0.3 | 10/30/2023 21:25 | 0.3 | 5.539 | 369687 | 0.2846 | 0.2846 | 94.9 | 5.539 | 112491 |
| Medium_QC_1 | QC | 0.5 | 10/30/2023 21:35 | 0.5 | 5.54 | 616000 | 0.4509 | 0.4509 | 90.2 | 5.54 | 119311 |
| High_QC_1 | QC | 0.8 | 10/30/2023 21:46 | 0.8 | 5.54 | 931149 | 0.753 | 0.753 | 94.1 | 5.54 | 108623 |
| preblank8 | DoubleBlank | | 10/30/2023 21:57 | | 5.574 | 1810 | 0.2364 | 0.2364 | | 5.562 | 660 |
| preblank9 | DoubleBlank | | 10/30/2023 22:07 | | 5.524 | 1053 | | | | | |
| LLOQ_2 | QC | 0.05 | 10/30/2023 22:18 | 0.05 | 5.545 | 73474 | 0.0463 | 0.0463 | 92.6 | 5.541 | 122893 |
| Low_QC_2 | QC | 0.3 | 10/30/2023 22:28 | 0.3 | 5.543 | 390134 | 0.3031 | 0.3031 | 101 | 5.539 | 111608 |
| Medium_QC_2 | QC | 0.5 | 10/30/2023 22:39 | 0.5 | 5.542 | 580124 | 0.4969 | 0.4969 | 99.4 | 5.542 | 102093 |
| High_QC_2 | QC | 0.8 | 10/30/2023 22:50 | 0.8 | 5.54 | 992912 | 0.7036 | 0.7036 | 87.9 | 5.54 | 123898 |
| preblank10 | DoubleBlank | | 10/30/2023 23:00 | | 5.778 | 5015 | 0.3617 | 0.3617 | | 5.698 | 1207 |
| preblank11 | DoubleBlank | | 10/30/2023 23:11 | | 5.553 | 1527 | 0.2053 | 0.2053 | | 5.566 | 638 |
| LLOQ_3 | QC | 0.05 | 10/30/2023 23:22 | 0.05 | 5.541 | 77083 | 0.0477 | 0.0477 | 95.4 | 5.541 | 125599 |
| Low_QC_3 | QC | 0.3 | 10/30/2023 23:32 | 0.3 | 5.539 | 386193 | 0.2637 | 0.2637 | 87.9 | 5.539 | 126575 |
| Medium_QC_3 | QC | 0.5 | 10/30/2023 23:43 | 0.5 | 5.539 | 595015 | 0.428 | 0.428 | 85.6 | 5.543 | 121329 |
| High_QC_3 | QC | 0.8 | 10/30/2023 23:54 | 0.8 | 5.54 | 958525 | 0.6946 | 0.6946 | 86.8 | 5.54 | 121129 |
| preblank12 | DoubleBlank | | 10/31/2023 0:04 | | 5.79 | 6191 | 0.5546 | 0.5546 | | 5.718 | 977 |
| preblank13 | DoubleBlank | | 10/31/2023 0:15 | | 5.792 | 2992 | 0.0854 | 0.0854 | | 5.603 | 2878 |
| LLOQ_4 | QC | 0.05 | 10/31/2023 0:26 | 0.05 | 5.541 | 71003 | 0.043 | 0.043 | 86.1 | 5.541 | 126578 |
| Low_QC_4 | QC | 0.3 | 10/31/2023 0:36 | 0.3 | 5.54 | 390058 | 0.2644 | 0.2644 | 88.1 | 5.54 | 127507 |
| Medium_QC_4 | QC | 0.5 | 10/31/2023 0:47 | 0.5 | 5.541 | 655047 | 0.4575 | 0.4575 | 91.5 | 5.541 | 125072 |
| High_QC_4 | QC | 0.8 | 10/31/2023 0:58 | 0.8 | 5.542 | 962721 | 0.6972 | 0.6972 | 87.1 | 5.542 | 121218 |
| preblank14 | DoubleBlank | | 10/31/2023 1:08 | | 5.536 | 1908 | 0.3901 | 0.3901 | | 5.595 | 426 |
| preblank15 | DoubleBlank | | 10/31/2023 1:19 | | 5.536 | 2087 | 0.9386 | 0.9386 | | 5.536 | 196 |
| LLOQ_5 | QC | 0.05 | 10/31/2023 1:30 | 0.05 | 5.543 | 74328 | 0.0439 | 0.0439 | 87.9 | 5.543 | 130140 |
| Low_QC_5 | QC | 0.3 | 10/31/2023 1:40 | 0.3 | 5.543 | 391296 | 0.2604 | 0.2604 | 86.8 | 5.543 | 129843 |
| Medium_QC_5 | QC | 0.5 | 10/31/2023 1:51 | 0.5 | 5.54 | 670410 | 0.4431 | 0.4431 | 88.6 | 5.54 | 132105 |
| High_QC_5 | QC | 0.8 | 10/31/2023 2:02 | 0.8 | 5.542 | 1005216 | 0.7028 | 0.7028 | 87.9 | 5.542 | 125562 |
| preblank16 | DoubleBlank | | 10/31/2023 2:12 | | 5.534 | 1902 | 0.1541 | 0.1541 | | 5.576 | 1049 |
| preblank17 | DoubleBlank | | 10/31/2023 2:23 | | 5.548 | 508 | 0.0205 | 0.0205 | | 5.611 | 1658 |
| LLOQ_6 | QC | 0.05 | 10/31/2023 2:33 | 0.05 | 5.544 | 74333 | 0.0439 | 0.0439 | 87.8 | 5.544 | 130229 |
| Low_QC_6 | QC | 0.3 | 10/31/2023 2:44 | 0.3 | 5.541 | 395976 | 0.2577 | 0.2577 | 85.9 | 5.541 | 132751 |
| Medium_QC_6 | QC | 0.5 | 10/31/2023 2:55 | 0.5 | 5.541 | 651874 | 0.4543 | 0.4543 | 90.9 | 5.537 | 125338 |
| High_QC_6 | QC | 0.8 | 10/31/2023 3:05 | 0.8 | 5.543 | 1014382 | 0.6735 | 0.6735 | 84.2 | 5.539 | 132169 |
| preblank18 | DoubleBlank | | 10/31/2023 3:16 | | 5.545 | 3000 | 0.4352 | 0.4352 | | 5.545 | 602 |
| preblank19 | DoubleBlank | | 10/31/2023 3:27 | | 5.632 | 554 | 0.0494 | 0.0494 | | 5.607 | 876 |
| zerocalibrator2 | Sample | | 10/31/2023 3:37 | | 5.562 | 1673 | 0 | 0 | | 5.537 | 132383 |
| L1_2 | Cal | 0.05 | 10/31/2023 3:48 | 0.05 | 5.542 | 84123 | 0.0495 | 0.0495 | 99 | 5.542 | 132724 |
| L2_2 | Cal | 0.08 | 10/31/2023 3:59 | 0.08 | 5.543 | 114528 | 0.0704 | 0.0704 | 88 | 5.543 | 131646 |
| L3_2 | Cal | 0.1 | 10/31/2023 4:09 | 0.1 | 5.543 | 153625 | 0.1034 | 0.1034 | 103.4 | 5.539 | 123707 |
| L4_2 | Cal | 0.15 | 10/31/2023 4:20 | 0.15 | 5.539 | 224553 | 0.1483 | 0.1483 | 98.9 | 5.539 | 128393 |
| L5_2 | Cal | 0.2 | 10/31/2023 4:31 | 0.2 | 5.538 | 295321 | 0.1985 | 0.1985 | 99.2 | 5.538 | 127587 |
| L6_2 | Cal | 0.3 | 10/31/2023 4:41 | 0.3 | 5.54 | 437193 | 0.2872 | 0.2872 | 95.7 | 5.54 | 131833 |
| L7_2 | Cal | 0.4 | 10/31/2023 4:52 | 0.4 | 5.539 | 568961 | 0.3719 | 0.3719 | 93 | 5.539 | 133191 |
| L8_2 | Cal | 0.5 | 10/31/2023 5:03 | 0.5 | 5.539 | 735176 | 0.5126 | 0.5126 | 102.5 | 5.539 | 125477 |
| L9_2 | Cal | 0.7 | 10/31/2023 5:13 | 0.7 | 5.539 | 983384 | 0.7039 | 0.7039 | 100.6 | 5.539 | 122646 |
| L10_2 | Cal | 0.8 | 10/31/2023 5:24 | 0.8 | 5.54 | 1086278 | 0.7378 | 0.7378 | 92.2 | 5.54 | 129322 |
| L11_2 | Cal | 0.9 | 10/31/2023 5:35 | 0.9 | 5.539 | 1260985 | 0.851 | 0.851 | 94.6 | 5.539 | 130307 |
| L12_2 | Cal | 1 | 10/31/2023 5:45 | 1 | 5.541 | 1346385 | 0.895 | 0.895 | 89.5 | 5.541 | 132342 |
| preblank20 | DoubleBlank | | 10/31/2023 5:56 | | 5.629 | 799 | 0.0393 | 0.0393 | | 5.633 | 1539 |
| **validation run 2** | | | | | | | | | | | |

| | | | | hydrazine Method | hydrazine Results | | | | | $^{15}N_2$-hydrazine (ISTD) Results | |
|---|---|---|---|---|---|---|---|---|---|---|---|
| Name | Type | Level | Acq. Date-Time | Exp. Conc. | RT | Resp. | Calc. Conc. | Final Conc. | Accuracy | RT | Resp. |
| preblank1 | DoubleBlank | | 11/7/2023 14:08 | | 5.796 | 1534 | 0.0237 | 0.0237 | | 5.641 | 4210 |
| preblank2 | DoubleBlank | | 11/7/2023 14:19 | | 5.344 | 269 | 0.0002 | 0.0002 | | 5.386 | 2923 |
| zerocalibrator1 | Blank | | 11/7/2023 14:29 | | | | | | | 5.543 | 123842 |
| L1_1 | Cal | 0.05 | 11/7/2023 14:40 | 0.05 | 5.543 | 86312 | 0.0504 | 0.0504 | 100.7 | 5.547 | 127935 |
| L2_1 | Cal | 0.08 | 11/7/2023 14:51 | 0.08 | 5.544 | 129028 | 0.0894 | 0.0894 | 111.8 | 5.544 | 114330 |
| L3_1 | Cal | 0.1 | 11/7/2023 15:01 | 0.1 | 5.544 | 163100 | 0.1101 | 0.1101 | 110.1 | 5.544 | 119227 |
| L4_1 | Cal | 0.15 | 11/7/2023 15:12 | 0.15 | 5.543 | 230221 | 0.1547 | 0.1547 | 103.2 | 5.543 | 121990 |
| L5_1 | Cal | 0.2 | 11/7/2023 15:23 | 0.2 | 5.547 | 321831 | 0.2164 | 0.2164 | 108.2 | 5.543 | 123628 |
| L6_1 | Cal | 0.3 | 11/7/2023 15:33 | 0.3 | 5.547 | 464358 | 0.3072 | 0.3072 | 102.4 | 5.547 | 126935 |
| L7_1 | Cal | 0.4 | 11/7/2023 15:44 | 0.4 | 5.545 | 637805 | 0.4402 | 0.4402 | 110 | 5.545 | 122585 |
| L8_1 | Cal | 0.5 | 11/7/2023 15:55 | 0.5 | 5.547 | 801139 | 0.561 | 0.561 | 112.2 | 5.543 | 121266 |
| L9_1 | Cal | 0.7 | 11/7/2023 16:05 | 0.7 | 5.548 | 1064502 | 0.7288 | 0.7288 | 104.1 | 5.544 | 124421 |
| L10_1 | Cal | 0.8 | 11/7/2023 16:16 | 0.8 | 5.548 | 1183735 | 0.821 | 0.821 | 102.6 | 5.544 | 122956 |
| L11_1 | Cal | 0.9 | 11/7/2023 16:26 | 0.9 | 5.547 | 1338678 | 0.9254 | 0.9254 | 102.8 | 5.547 | 123494 |
| L12_1 | Cal | 1 | 11/7/2023 16:37 | 1 | 5.545 | 1529572 | 0.9691 | 0.9691 | 96.9 | 5.545 | 134797 |
| preblank3 | DoubleBlank | | 11/7/2023 16:48 | | 5.707 | 306 | 0.0331 | 0.0331 | | 5.64 | 645 |
| preblank4 | DoubleBlank | | 11/7/2023 16:58 | | 5.538 | 988 | 0.0113 | 0.0113 | | 5.564 | 4464 |
| LLOQ_1 | QC | 0.05 | 11/7/2023 17:09 | 0.05 | 5.547 | 78666 | 0.0474 | 0.0474 | 94.8 | 5.547 | 122871 |
| Low_QC_1 | QC | 0.3 | 11/7/2023 17:20 | 0.3 | 5.546 | 491442 | 0.3319 | 0.3319 | 110.6 | 5.546 | 124554 |
| Medium_QC_1 | QC | 0.5 | 11/7/2023 17:30 | 0.5 | 5.544 | 748987 | 0.5139 | 0.5139 | 102.8 | 5.544 | 123620 |
| High_QC_1 | QC | 0.8 | 11/7/2023 17:41 | 0.8 | 5.545 | 1139890 | 0.8603 | 0.8603 | 107.5 | 5.545 | 113041 |
| preblank5 | DoubleBlank | | 11/7/2023 17:52 | | 5.552 | 3211 | 0.7121 | 0.7121 | | 5.544 | 384 |
| preblank6 | DoubleBlank | | 11/7/2023 18:02 | | 5.548 | 6052 | | | | | |
| LLOQ_2 | QC | 0.05 | 11/7/2023 18:13 | 0.05 | 5.547 | 85051 | 0.0499 | 0.0499 | 99.7 | 5.543 | 127180 |
| Low_QC_2 | QC | 0.3 | 11/7/2023 18:24 | 0.3 | 5.544 | 483068 | 0.3336 | 0.3336 | 111.2 | 5.544 | 121832 |
| Medium_QC_2 | QC | 0.5 | 11/7/2023 18:34 | 0.5 | 5.545 | 743634 | 0.5114 | 0.5114 | 102.3 | 5.545 | 123322 |
| High_QC_2 | QC | 0.8 | 11/7/2023 18:45 | 0.8 | 5.546 | 1175630 | 0.7873 | 0.7873 | 98.4 | 5.546 | 127295 |
| preblank7 | DoubleBlank | | 11/7/2023 18:56 | | 5.55 | 1955 | 0.5488 | 0.5488 | | 5.6 | 302 |
| preblank118 | DoubleBlank | | 11/7/2023 19:06 | | 5.702 | 275 | 0.0095 | 0.0095 | | 5.719 | 1378 |
| LLOQ_3 | QC | 0.05 | 11/7/2023 19:17 | 0.05 | 5.545 | 87483 | 0.0545 | 0.0545 | 109.1 | 5.545 | 120969 |
| Low_QC_3 | QC | 0.3 | 11/7/2023 19:27 | 0.3 | 5.545 | 482058 | 0.3301 | 0.3301 | 110 | 5.545 | 122826 |
| Medium_QC_3 | QC | 0.5 | 11/7/2023 19:38 | 0.5 | 5.548 | 21079036 | 20.3802 | 20.3802 | 4076 | 5.547 | 89001 |
| High_QC_3 | QC | 0.8 | 11/7/2023 19:49 | 0.8 | 5.546 | 1157850 | 0.819 | 0.819 | 102.4 | 5.546 | 120559 |
| preblank9 | DoubleBlank | | 11/7/2023 19:59 | | 5.571 | 1455 | 0.0846 | 0.0846 | | 5.659 | 1357 |
| preblank10 | DoubleBlank | | 11/7/2023 20:10 | | 5.732 | 104 | | | | | |
| LLOQ_4 | QC | 0.05 | 11/7/2023 20:21 | 0.05 | 5.544 | 92779 | 0.0512 | 0.0512 | 102.3 | 5.544 | 135691 |
| Low_QC_4 | QC | 0.3 | 11/7/2023 20:31 | 0.3 | 5.545 | 507887 | 0.331 | 0.331 | 110.3 | 5.545 | 129095 |
| Medium_QC_4 | QC | 0.5 | 11/7/2023 20:42 | 0.5 | 5.546 | 741842 | 0.5134 | 0.5134 | 102.7 | 5.546 | 122536 |
| High_QC_4 | QC | 0.8 | 11/7/2023 20:53 | 0.8 | 5.545 | 1110019 | 0.7056 | 0.7056 | 88.2 | 5.545 | 133965 |
| preblank11 | DoubleBlank | | 11/7/2023 21:03 | | 5.773 | 3322 | 0.9824 | 0.9824 | | 5.731 | 289 |
| preblank12 | DoubleBlank | | 11/7/2023 21:14 | | 5.791 | 4487 | 0.1178 | 0.1178 | | 5.736 | 3078 |
| LLOQ_5 | QC | 0.05 | 11/7/2023 21:25 | 0.05 | 5.547 | 95329 | 0.0501 | 0.0501 | 100.2 | 5.547 | 141971 |
| Low_QC_5 | QC | 0.3 | 11/7/2023 21:35 | 0.3 | 5.545 | 474849 | 0.3185 | 0.3185 | 106.2 | 5.545 | 125319 |
| Medium_QC_5 | QC | 0.5 | 11/7/2023 21:46 | 0.5 | 5.545 | 754737 | 0.4898 | 0.4898 | 98 | 5.545 | 130579 |
| High_QC_5 | QC | 0.8 | 11/7/2023 21:57 | 0.8 | 5.544 | 1147181 | 0.769 | 0.769 | 96.1 | 5.544 | 127138 |
| preblank13 | DoubleBlank | | 11/7/2023 22:07 | | 5.479 | 572 | 0.7794 | 0.7794 | | 5.504 | 63 |
| preblank14 | DoubleBlank | | 11/7/2023 22:18 | | 5.55 | 18977 | 1.3205 | 1.3205 | | 5.525 | 1230 |
| LLOQ_6 | QC | 0.05 | 11/7/2023 22:29 | 0.05 | 5.542 | 91128 | 0.0526 | 0.0526 | 105.2 | 5.546 | 130093 |
| Low_QC_6 | QC | 0.3 | 11/7/2023 22:39 | 0.3 | 5.545 | 510365 | 0.3297 | 0.3297 | 109.9 | 5.541 | 130201 |
| Medium_QC_6 | QC | 0.5 | 11/7/2023 22:50 | 0.5 | 5.546 | 730888 | 0.5176 | 0.5176 | 103.5 | 5.546 | 119764 |
| High_QC_6 | QC | 0.8 | 11/7/2023 23:00 | 0.8 | 5.545 | 1194901 | 0.73 | 0.73 | 91.2 | 5.545 | 139437 |
| preblank15 | DoubleBlank | | 11/7/2023 23:11 | | 5.542 | 114742 | 18.4279 | 18.4279 | | 5.521 | 536 |
| preblank16 | DoubleBlank | | 11/7/2023 23:22 | | 5.779 | 3887 | 2.8172 | 2.8172 | | 5.733 | 118 |
| stab_stock_L1_-80C | Sample | 0.3 | 11/7/2023 23:32 | 0.3 | 5.544 | 487855 | 0.2844 | 0.2844 | 94.8 | 5.544 | 143758 |
| stab_stock_H1_-80C | Sample | 0.8 | 11/7/2023 23:43 | 0.8 | 5.547 | 1142770 | 0.8044 | 0.8044 | 100.5 | 5.543 | 121139 |
| preblank17 | DoubleBlank | | 11/7/2023 23:54 | | 5.545 | 103281 | 62.617 | 62.617 | | 5.612 | 142 |
| preblank18 | DoubleBlank | | 11/8/2023 0:05 | | 5.545 | 71992 | 37.6667 | 37.6667 | | 5.377 | 164 |
| stab_stock_L1_-20C | Sample | 0.3 | 11/8/2023 0:16 | 0.3 | 5.546 | 388834 | 0.2447 | 0.2447 | 81.6 | 5.546 | 132619 |
| stab_stock_H1_-20C | Sample | 0.8 | 11/8/2023 0:27 | 0.8 | 5.547 | 1028113 | 0.6544 | 0.6544 | 81.8 | 5.547 | 133659 |
| preblank19 | DoubleBlank | | 11/8/2023 0:37 | | 5.548 | 112045 | 6.06 | 6.06 | | 5.552 | 1590 |
| preblank20 | DoubleBlank | | 11/8/2023 0:48 | | 5.55 | 72121 | 11.4142 | 11.4142 | | 5.382 | 544 |
| stab_stock_L1_4C | Sample | 0.3 | 11/8/2023 0:59 | 0.3 | 5.547 | 518707 | 0.3012 | 0.3012 | 100.4 | 5.547 | 144533 |
| stab_stock_H1_4C | Sample | 0.8 | 11/8/2023 1:10 | 0.8 | 5.543 | 1262007 | 0.7721 | 0.7721 | 96.5 | 5.543 | 139309 |
| preblank21 | DoubleBlank | | 11/8/2023 1:20 | | 5.543 | 65446 | 5.4783 | 5.4783 | | 5.581 | 1027 |
| preblank22 | DoubleBlank | | 11/8/2023 1:31 | | 5.548 | 66920 | | | | | |
| stab_stock_L1_auto | Sample | 0.3 | 11/8/2023 1:42 | 0.3 | 5.548 | 462225 | 0.2924 | 0.2924 | 97.5 | 5.548 | 132599 |
| stab_stock_H1_auto | Sample | 0.8 | 11/8/2023 1:53 | 0.8 | 5.547 | 1285336 | 0.7979 | 0.7979 | 99.7 | 5.547 | 137336 |
| preblank23 | DoubleBlank | | 11/8/2023 2:04 | | 5.778 | 2818 | 4.0312 | 4.0312 | | 5.748 | 60 |
| preblank24 | DoubleBlank | | 11/8/2023 2:15 | | 5.788 | 2935 | 0.1398 | 0.1398 | | 5.683 | 1712 |
| stab_stock_L1_RT | Sample | 0.3 | 11/8/2023 2:25 | 0.3 | 5.544 | 405424 | 0.2605 | 0.2605 | 86.8 | 5.544 | 130143 |
| stab_stock_H1_RT | Sample | 0.8 | 11/8/2023 2:36 | 0.8 | 5.546 | 1142231 | 0.7572 | 0.7572 | 94.7 | 5.546 | 128544 |
| preblank25 | DoubleBlank | | 11/8/2023 2:47 | | 5.547 | 49084 | | | | | |
| preblank26 | DoubleBlank | | 11/8/2023 2:58 | | 5.545 | 38141 | 3.8294 | 3.8294 | | 5.6 | 856 |
| stab_stock_L2_-80C | Sample | 0.3 | 11/8/2023 3:08 | 0.3 | 5.547 | 504139 | 0.316 | 0.316 | 105.3 | 5.547 | 134066 |
| stab_stock_H2_-80C | Sample | 0.8 | 11/8/2023 3:19 | 0.8 | 5.545 | 1156747 | 0.8023 | 0.8023 | 100.3 | 5.545 | 122928 |
| preblank27 | DoubleBlank | | 11/8/2023 3:30 | | 5.547 | 43958 | 2.361 | 2.361 | | 5.572 | 1597 |
| preblank28 | DoubleBlank | | 11/8/2023 3:41 | | 5.548 | 37960 | 2.6434 | 2.6434 | | 5.662 | 1233 |
| stab_stock_L2_-20C | Sample | 0.3 | 11/8/2023 3:51 | 0.3 | 5.546 | 376270 | 0.2495 | 0.2495 | 83.2 | 5.546 | 125917 |
| stab_stock_H2_-20C | Sample | 0.8 | 11/8/2023 4:02 | 0.8 | 5.543 | 1260666 | 0.7605 | 0.7605 | 95.1 | 5.543 | 141269 |
| preblank29 | DoubleBlank | | 11/8/2023 4:13 | | 5.545 | 31308 | 4.87 | 4.87 | | 5.386 | 553 |
| preblank30 | DoubleBlank | | 11/8/2023 4:24 | | 5.787 | 2913 | 2.0583 | 2.0583 | | 5.74 | 121 |
| stab_stock_L2_4C | Sample | 0.3 | 11/8/2023 4:35 | 0.3 | 5.546 | 501128 | 0.3004 | 0.3004 | 100.1 | 5.542 | 139998 |
| stab_stock_H2_4C | Sample | 0.8 | 11/8/2023 4:46 | 0.8 | 5.546 | 1295208 | 0.8225 | 0.8225 | 102.8 | 5.546 | 134296 |
| preblank31 | DoubleBlank | | 11/8/2023 4:56 | | 5.547 | 41629 | 2.1558 | 2.1558 | | 5.573 | 1656 |
| preblank32 | DoubleBlank | | 11/8/2023 5:07 | | 5.546 | 40040 | | | | | |

| | | | | hydrazine Method | hydrazine Results | | | | | $^{15}N_2$-hydrazine (ISTD) Results | |
|---|---|---|---|---|---|---|---|---|---|---|---|
| Name | Type | Level | Acq. Date-Time | Exp. Conc. | RT | Resp. | Calc. Conc. | Final Conc. | Accuracy | RT | Resp. |
| stab_stock_L2_auto | Sample | 0.3 | 11/8/2023 5:18 | 0.3 | 5.548 | 487720 | 0.3098 | 0.3098 | 103.3 | 5.543 | 132248 |
| stab_stock_H2_auto | Sample | 0.8 | 11/8/2023 5:29 | 0.8 | 5.545 | 1267091 | 0.7967 | 0.7967 | 99.6 | 5.545 | 135592 |
| preblank33 | DoubleBlank | | 11/8/2023 5:39 | | 5.542 | 40188 | 0.5786 | 0.5786 | | 5.605 | 5900 |
| preblank34 | DoubleBlank | | 11/8/2023 5:50 | | 5.544 | 41247 | 8.8112 | 8.8112 | | 5.569 | 403 |
| stab_stock_L2_RT | Sample | 0.3 | 11/8/2023 6:01 | 0.3 | 5.545 | 419676 | 0.2497 | 0.2497 | 83.2 | 5.545 | 140353 |
| stab_stock_H2_RT | Sample | 0.8 | 11/8/2023 6:12 | 0.8 | 5.544 | 1144923 | 0.7226 | 0.7226 | 90.3 | 5.544 | 134964 |
| preblank35 | DoubleBlank | | 11/8/2023 6:23 | | 5.543 | 38854 | 2.2075 | 2.2075 | | 5.618 | 1510 |
| preblank36 | DoubleBlank | | 11/8/2023 6:33 | | 5.79 | 3435 | 0.0379 | 0.0379 | | 5.69 | 6491 |
| stab_stock_L3_-80C | Sample | 0.3 | 11/8/2023 6:44 | 0.3 | 5.545 | 464337 | 0.2845 | 0.2845 | 94.8 | 5.545 | 136796 |
| stab_stock_H3_-80C | Sample | 0.8 | 11/8/2023 6:55 | 0.8 | 5.547 | 1241211 | 0.7667 | 0.7667 | 95.8 | 5.542 | 137974 |
| preblank37 | DoubleBlank | | 11/8/2023 7:06 | | 5.545 | 38862 | | | | | |
| preblank38 | DoubleBlank | | 11/8/2023 7:17 | | 5.789 | 2744 | 2.6045 | 2.6045 | | 5.735 | 90 |
| stab_stock_L3_-20C | Sample | 0.3 | 11/8/2023 7:27 | 0.3 | 5.543 | 392441 | 0.2344 | 0.2344 | 78.1 | 5.543 | 139544 |
| stab_stock_H3_-20C | Sample | 0.8 | 11/8/2023 7:38 | 0.8 | 5.544 | 1172462 | 0.7238 | 0.7238 | 90.5 | 5.544 | 137972 |
| preblank39 | DoubleBlank | | 11/8/2023 7:49 | | 5.542 | 39567 | 8.2372 | 8.2372 | | 5.593 | 413 |
| preblank40 | DoubleBlank | | 11/8/2023 8:00 | | 5.547 | 45118 | | | | | |
| stab_stock_L3_4C | Sample | 0.3 | 11/8/2023 8:10 | 0.3 | 5.544 | 483291 | 0.3061 | 0.3061 | 102 | 5.544 | 132567 |
| stab_stock_H3_4C | Sample | 0.8 | 11/8/2023 8:21 | 0.8 | 5.546 | 1353847 | 0.8329 | 0.8329 | 104.1 | 5.546 | 138637 |
| preblank41 | DoubleBlank | | 11/8/2023 8:32 | | 5.544 | 38516 | 4.8416 | 4.8416 | | 5.607 | 684 |
| preblank42 | DoubleBlank | | 11/8/2023 8:43 | | 5.546 | 36928 | 2.1889 | 2.1889 | | 5.656 | 1447 |
| stab_stock_L3_auto | Sample | 0.3 | 11/8/2023 8:53 | 0.3 | 5.547 | 488588 | 0.3038 | 0.3038 | 101.3 | 5.547 | 135015 |
| stab_stock_H3_auto | Sample | 0.8 | 11/8/2023 9:04 | 0.8 | 5.547 | 1349771 | 0.8162 | 0.8162 | 102 | 5.543 | 141021 |
| preblank43 | DoubleBlank | | 11/8/2023 9:15 | | 5.545 | 36653 | 5.5896 | 5.5896 | | 5.637 | 564 |
| preblank44 | DoubleBlank | | 11/8/2023 9:26 | | 5.552 | 34083 | 9.6102 | 9.6102 | | 5.384 | 305 |
| stab_stock_L3_RT | Sample | 0.3 | 11/8/2023 9:37 | 0.3 | 5.544 | 368220 | 0.2201 | 0.2201 | 73.4 | 5.544 | 139119 |
| stab_stock_H3_RT | Sample | 0.8 | 11/8/2023 9:48 | 0.8 | 5.543 | 1184698 | 0.6932 | 0.6932 | 86.6 | 5.543 | 145507 |
| preblank45 | DoubleBlank | | 11/8/2023 9:58 | | 5.544 | 37521 | 7.1663 | 7.1663 | | 5.511 | 450 |
| preblank46 | DoubleBlank | | 11/8/2023 10:09 | | 5.544 | 37996 | 0.9729 | 0.9729 | | 5.603 | 3335 |
| stab_dev_L1_-80C | Sample | 0.3 | 11/8/2023 10:20 | 0.3 | 5.546 | 441387 | 0.2781 | 0.2781 | 92.7 | 5.546 | 132962 |
| stab_dev_H1_-80C | Sample | 0.8 | 11/8/2023 10:31 | 0.8 | 5.543 | 1176531 | 0.8075 | 0.8075 | 100.9 | 5.543 | 124231 |
| preblank47 | DoubleBlank | | 11/8/2023 10:41 | | 5.778 | 2060 | 0.035 | 0.035 | | 5.727 | 4154 |
| preblank48 | DoubleBlank | | 11/8/2023 10:52 | | | | | | | 5.741 | 215 |
| stab_dev_L1_-20C | Sample | 0.3 | 11/8/2023 11:03 | 0.3 | 5.547 | 356266 | 0.1981 | 0.1981 | 66 | 5.543 | 148998 |
| stab_dev_H1_-20C | Sample | 0.8 | 11/8/2023 11:14 | 0.8 | 5.548 | 938106 | 0.5368 | 0.5368 | 67.1 | 5.548 | 148311 |
| preblank49 | DoubleBlank | | 11/8/2023 11:25 | | | | | | | 5.73 | 4731 |
| preblank50 | DoubleBlank | | 11/8/2023 11:35 | | 5.545 | 36586 | 1.0243 | 1.0243 | | 5.591 | 3052 |
| stab_dev_L1_4C | Sample | 0.3 | 11/8/2023 11:46 | 0.3 | 5.546 | 309903 | 0.2008 | 0.2008 | 66.9 | 5.546 | 127964 |
| stab_dev_H1_4C | Sample | 0.8 | 11/8/2023 11:57 | 0.8 | 5.545 | 978374 | 0.5521 | 0.5521 | 69 | 5.545 | 150436 |
| preblank51_ | DoubleBlank | | 11/8/2023 12:08 | | 5.535 | 4832 | 0.4013 | 0.4013 | | 5.568 | 1017 |
| preblank52_ | DoubleBlank | | 11/8/2023 12:19 | | 5.378 | 820 | 1.2365 | 1.2365 | | 5.383 | 57 |
| stab_dev_L1_auto | Sample | 0.3 | 11/8/2023 12:29 | 0.3 | 5.543 | 451874 | 0.2579 | 0.2579 | 86 | 5.543 | 146459 |
| stab_dev_H1_auto | Sample | 0.8 | 11/8/2023 12:40 | 0.8 | 5.547 | 1138764 | 0.7052 | 0.7052 | 88.1 | 5.547 | 137507 |
| preblank53_ | DoubleBlank | | 11/8/2023 12:51 | | 5.535 | 1733 | 0.0144 | 0.0144 | | 5.652 | 6762 |
| preblank54_ | DoubleBlank | | 11/8/2023 13:02 | | 5.542 | 1719 | 0.1029 | 0.1029 | | 5.643 | 1338 |
| stab_dev_L1_RT | Sample | 0.3 | 11/8/2023 13:12 | 0.3 | 5.546 | 388433 | 0.2485 | 0.2485 | 82.8 | 5.542 | 130495 |
| stab_dev_H1_RT | Sample | 0.8 | 11/8/2023 13:23 | 0.8 | 5.544 | 1071902 | 0.6641 | 0.6641 | 83 | 5.544 | 137354 |
| preblank55_ | DoubleBlank | | 11/8/2023 13:34 | | 5.643 | 609 | 0.0452 | 0.0452 | | 5.63 | 991 |
| preblank56_ | DoubleBlank | | 11/8/2023 13:45 | | 5.577 | 1444 | 0.0247 | 0.0247 | | 5.61 | 3839 |
| stab_dev_L2_-80C | Sample | 0.3 | 11/8/2023 13:55 | 0.3 | 5.547 | 436769 | 0.2665 | 0.2665 | 88.8 | 5.543 | 137111 |
| stab_dev_H2_-80C | Sample | 0.8 | 11/8/2023 14:06 | 0.8 | 5.545 | 1120409 | 0.6959 | 0.6959 | 87 | 5.545 | 137083 |
| preblank57_ | DoubleBlank | | 11/8/2023 14:17 | | 5.617 | 727 | 0.0023 | 0.0023 | | 5.621 | 6249 |
| preblank58_ | DoubleBlank | | 11/8/2023 14:28 | | 5.542 | 5357 | 0.2283 | 0.2283 | | 5.508 | 1954 |
| stab_dev_L2_-20C | Sample | 0.3 | 11/8/2023 14:39 | 0.3 | 5.545 | 359314 | 0.2104 | 0.2104 | 70.1 | 5.545 | 141824 |
| stab_dev_H2_-20C | Sample | 0.8 | 11/8/2023 14:49 | 0.8 | 5.544 | 1056439 | 0.6479 | 0.6479 | 81 | 5.544 | 138706 |
| preblank59_ | DoubleBlank | | 11/8/2023 15:00 | | 5.563 | 10876 | 4.3833 | 4.3833 | | 5.554 | 213 |
| preblank60_ | DoubleBlank | | 11/8/2023 15:11 | | 5.545 | 1808 | 0.0618 | 0.0618 | | 5.575 | 2239 |
| stab_dev_L2_4C | Sample | 0.3 | 11/8/2023 15:22 | 0.3 | 5.543 | 344469 | 0.2063 | 0.2063 | 68.8 | 5.548 | 138535 |
| stab_dev_H2_4C | Sample | 0.8 | 11/8/2023 15:33 | 0.8 | 5.545 | 1057067 | 0.6609 | 0.6609 | 82.6 | 5.545 | 136106 |
| preblank61_ | DoubleBlank | | 11/8/2023 15:43 | | 5.337 | 188 | 0.0085 | 0.0085 | | 5.522 | 996 |
| preblank62_ | DoubleBlank | | 11/8/2023 15:54 | | 5.799 | 3675 | 0.4451 | 0.4451 | | 5.698 | 699 |
| stab_dev_L2_auto | Sample | 0.3 | 11/8/2023 16:05 | 0.3 | 5.544 | 458065 | 0.2772 | 0.2772 | 92.4 | 5.544 | 138399 |
| stab_dev_H2_auto | Sample | 0.8 | 11/8/2023 16:16 | 0.8 | 5.546 | 1126796 | 0.7144 | 0.7144 | 89.3 | 5.542 | 134334 |
| preblank63_ | DoubleBlank | | 11/8/2023 16:26 | | 5.544 | 2885 | 0.3608 | 0.3608 | | 5.54 | 674 |
| preblank64_ | DoubleBlank | | 11/8/2023 16:37 | | 5.553 | 878 | 0.1001 | 0.1001 | | 5.494 | 701 |
| stab_dev_L2_RT | Sample | 0.3 | 11/8/2023 16:48 | 0.3 | 5.545 | 417057 | 0.2712 | 0.2712 | 90.4 | 5.545 | 128716 |
| stab_dev_H2_RT | Sample | 0.8 | 11/8/2023 16:59 | 0.8 | 5.545 | 1058146 | 0.6926 | 0.6926 | 86.6 | 5.545 | 130067 |
| preblank65_ | DoubleBlank | | 11/8/2023 17:10 | | 5.546 | 2535 | 0.108 | 0.108 | | 5.575 | 1886 |
| preblank66_ | DoubleBlank | | 11/8/2023 17:20 | | 5.548 | 1320 | | | | | |
| stab_dev_L3_-80C | Sample | 0.3 | 11/8/2023 17:31 | 0.3 | 5.546 | 427288 | 0.2557 | 0.2557 | 85.2 | 5.542 | 139647 |
| stab_dev_H3_-80C | Sample | 0.8 | 11/8/2023 17:42 | 0.8 | 5.545 | 1015005 | 0.712 | 0.712 | 89 | 5.545 | 121410 |
| preblank67_ | DoubleBlank | | 11/8/2023 17:53 | | 5.531 | 5392 | | | | | |
| preblank78_ | DoubleBlank | | 11/8/2023 18:04 | | 5.552 | 4081 | 0.2684 | 0.2684 | | 5.557 | 1272 |
| stab_dev_L3_-20C | Sample | 0.3 | 11/8/2023 18:14 | 0.3 | 5.546 | 256492 | 0.1579 | 0.1579 | 52.6 | 5.546 | 133351 |
| stab_dev_H3_-20C | Sample | 0.8 | 11/8/2023 18:25 | 0.8 | 5.545 | 874883 | 0.5689 | 0.5689 | 71.1 | 5.545 | 130613 |
| preblank69_ | DoubleBlank | | 11/8/2023 18:36 | | 5.401 | 107 | 0 | 0 | | 5.582 | 2676 |
| preblank70_ | DoubleBlank | | 11/8/2023 18:47 | | 5.783 | 2713 | 0.9736 | 0.9736 | | 5.72 | 238 |
| stab_dev_L3_4C | Sample | 0.3 | 11/8/2023 18:57 | 0.3 | 5.544 | 343675 | 0.1838 | 0.1838 | 61.3 | 5.544 | 154482 |
| stab_dev_H3_4C | Sample | 0.8 | 11/8/2023 19:08 | 0.8 | 5.547 | 989948 | 0.5615 | 0.5615 | 70.2 | 5.542 | 149723 |
| preblank71_ | DoubleBlank | | 11/8/2023 19:19 | | 5.557 | 1462 | 0.0122 | 0.0122 | | 5.612 | 6311 |
| preblank72_ | DoubleBlank | | 11/8/2023 19:30 | | 5.704 | 153 | 0.0015 | 0.0015 | | 5.524 | 1430 |
| stab_dev_L3_auto | Sample | 0.3 | 11/8/2023 19:41 | 0.3 | 5.544 | 426392 | 0.2628 | 0.2628 | 87.6 | 5.544 | 135693 |
| stab_dev_H3_auto | Sample | 0.8 | 11/8/2023 19:51 | 0.8 | 5.547 | 1144084 | 0.6969 | 0.6969 | 87.1 | 5.543 | 139776 |
| preblank73_ | DoubleBlank | | 11/8/2023 20:02 | | | | | | | 5.558 | 339 |
| preblank74_ | DoubleBlank | | 11/8/2023 20:13 | | | | | | | 5.532 | 225 |
| stab_dev_L3_RT | Sample | 0.3 | 11/8/2023 20:24 | 0.3 | 5.546 | 415558 | 0.2539 | 0.2539 | 84.6 | 5.542 | 136730 |

| | | | | hydrazine Method | hydrazine Results | | | | | $^{15}N_2$-hydrazine (ISTD) Results | |
|---|---|---|---|---|---|---|---|---|---|---|---|
| **Name** | **Type** | **Level** | **Acq. Date-Time** | **Exp. Conc.** | **RT** | **Resp.** | **Calc. Conc.** | **Final Conc.** | **Accuracy** | **RT** | **Resp.** |
| stab_dev_H3_RT | Sample | 0.8 | 11/8/2023 20:35 | 0.8 | 5.545 | 1044501 | 0.6814 | 0.6814 | 85.2 | 5.545 | 130486 |
| preblank75_ | DoubleBlank | | 11/8/2023 20:45 | | 5.539 | 1520 | 0.0389 | 0.0389 | | 5.577 | 2807 |
| preblank76_ | DoubleBlank | | 11/8/2023 20:56 | | 5.55 | 3026 | | | | | |
| zerocalibrator2 | Blank | | 11/8/2023 21:07 | | 5.561 | 1216 | 0 | 0 | | 5.544 | 135666 |
| L1_2 | Cal | 0.05 | 11/8/2023 21:18 | 0.05 | 5.549 | 79357 | 0.0431 | 0.0431 | 86.2 | 5.545 | 134510 |
| L2_2 | Cal | 0.08 | 11/8/2023 21:28 | 0.08 | 5.549 | 133009 | 0.0785 | 0.0785 | 98.1 | 5.544 | 132886 |
| L3_2 | Cal | 0.1 | 11/8/2023 21:39 | 0.1 | 5.542 | 164346 | 0.1065 | 0.1065 | 106.5 | 5.542 | 123880 |
| L4_2 | Cal | 0.15 | 11/8/2023 21:50 | 0.15 | 5.546 | 233815 | 0.1424 | 0.1424 | 95 | 5.546 | 134062 |
| L5_2 | Cal | 0.2 | 11/8/2023 22:00 | 0.2 | 5.542 | 303328 | 0.186 | 0.186 | 93 | 5.542 | 134802 |
| L6_2 | Cal | 0.3 | 11/8/2023 22:11 | 0.3 | 5.545 | 454581 | 0.2804 | 0.2804 | 93.5 | 5.545 | 135819 |
| L7_2 | Cal | 0.4 | 11/8/2023 22:21 | 0.4 | 5.544 | 586598 | 0.3528 | 0.3528 | 88.2 | 5.544 | 140081 |
| L8_2 | Cal | 0.5 | 11/8/2023 22:32 | 0.5 | 5.547 | 740661 | 0.4648 | 0.4648 | 93 | 5.547 | 134947 |
| L9_2 | Cal | 0.7 | 11/8/2023 22:43 | 0.7 | 5.546 | 1030965 | 0.6798 | 0.6798 | 97.1 | 5.546 | 129088 |
| L10_2 | Cal | 0.8 | 11/8/2023 22:53 | 0.8 | 5.543 | 1161300 | 0.7651 | 0.7651 | 95.6 | 5.543 | 129359 |
| L11_2 | Cal | 0.9 | 11/8/2023 23:04 | 0.9 | 5.543 | 1272028 | 0.8353 | 0.8353 | 92.8 | 5.543 | 129888 |
| L12_2 | Cal | 1 | 11/8/2023 23:15 | 1 | 5.544 | 1395824 | 0.9608 | 0.9608 | 96.1 | 5.544 | 124070 |
| preblank77_ | DoubleBlank | | 11/8/2023 23:25 | | 5.548 | 4061 | 0.0662 | 0.0662 | | 5.565 | 4729 |
| preblank78_ | DoubleBlank | | 11/8/2023 23:36 | | 5.55 | 3631 | | | | | |
| **validation run 3** | | | | | | | | | | | |
| preblank1 | DoubleBlank | | 11/14/2023 11:42 | | 5.542 | 250003 | 11.1766 | 11.1766 | | 5.63 | 2078 |
| preblank2 | DoubleBlank | | 11/14/2023 11:53 | | 5.544 | 285819 | 34.7905 | 34.7905 | | 5.539 | 764 |
| zerocalibrator1 | Blank | | 11/14/2023 12:03 | | 5.723 | 463 | 0 | 0 | | 5.542 | 136465 |
| L1_1 | Cal | 0.05 | 11/14/2023 12:14 | 0.05 | 5.543 | 94008 | 0.0478 | 0.0478 | 95.6 | 5.543 | 140159 |
| L2_1 | Cal | 0.08 | 11/14/2023 12:25 | 0.08 | 5.544 | 147699 | 0.0869 | 0.0869 | 108.6 | 5.544 | 135429 |
| L3_1 | Cal | 0.1 | 11/14/2023 12:35 | 0.1 | 5.545 | 177161 | 0.1006 | 0.1006 | 100.6 | 5.545 | 143126 |
| L4_1 | Cal | 0.15 | 11/14/2023 12:46 | 0.15 | 5.544 | 258720 | 0.1641 | 0.1641 | 109.4 | 5.544 | 134729 |
| L5_1 | Cal | 0.2 | 11/14/2023 12:57 | 0.2 | 5.548 | 320131 | 0.1883 | 0.1883 | 94.2 | 5.544 | 146781 |
| L6_1 | Cal | 0.3 | 11/14/2023 13:07 | 0.3 | 5.544 | 490097 | 0.3139 | 0.3139 | 104.6 | 5.544 | 138821 |
| L7_1 | Cal | 0.4 | 11/14/2023 13:18 | 0.4 | 5.547 | 652026 | 0.4047 | 0.4047 | 101.2 | 5.547 | 144687 |
| L8_1 | Cal | 0.5 | 11/14/2023 13:29 | 0.5 | 5.544 | 800935 | 0.4996 | 0.4996 | 99.9 | 5.544 | 144922 |
| L9_1 | Cal | 0.7 | 11/14/2023 13:39 | 0.7 | 5.545 | 1101478 | 0.7432 | 0.7432 | 106.2 | 5.545 | 135227 |
| L10_1 | Cal | 0.8 | 11/14/2023 13:50 | 0.8 | 5.544 | 1226111 | 0.8153 | 0.8153 | 101.9 | 5.544 | 137455 |
| L11_1 | Cal | 0.9 | 11/14/2023 14:01 | 0.9 | 5.544 | 1375358 | 0.9005 | 0.9005 | 100.1 | 5.544 | 139825 |
| L12_1 | Cal | 1 | 11/14/2023 14:11 | 1 | 5.545 | 1494952 | 0.993 | 0.993 | 99.3 | 5.545 | 138029 |
| preblank3 | DoubleBlank | | 11/14/2023 14:22 | | 5.546 | 272195 | 39.6525 | 39.6525 | | 5.554 | 638 |
| preblank4 | DoubleBlank | | 11/14/2023 14:32 | | 5.542 | 247803 | 72.6136 | 72.6136 | | 5.374 | 317 |
| LLOQ_1 | QC | 0.05 | 11/14/2023 14:43 | 0.05 | 5.544 | 104884 | 0.052 | 0.052 | 103.9 | 5.544 | 146659 |
| Low_QC_1 | QC | 0.3 | 11/14/2023 14:54 | 0.3 | 5.544 | 556823 | 0.3634 | 0.3634 | 121.1 | 5.544 | 137049 |
| Medium_QC_1 | QC | 0.5 | 11/14/2023 15:04 | 0.5 | 5.546 | 827108 | 0.547 | 0.547 | 109.4 | 5.542 | 137017 |
| High_QC_1 | QC | 0.8 | 11/14/2023 15:15 | 0.8 | 5.544 | 1283179 | 0.8244 | 0.8244 | 103.1 | 5.544 | 142280 |
| preblank5 | DoubleBlank | | 11/14/2023 15:26 | | 5.546 | 301441 | 12.9354 | 12.9354 | | 5.567 | 2166 |
| preblank6 | DoubleBlank | | 11/14/2023 15:36 | | 5.543 | 313017 | 13.4931 | 13.4931 | | 5.551 | 2156 |
| LLOQ_2 | QC | 0.05 | 11/14/2023 15:47 | 0.05 | 5.542 | 97483 | 0.0504 | 0.0504 | 100.7 | 5.546 | 139648 |
| Low_QC_2 | QC | 0.3 | 11/14/2023 15:58 | 0.3 | 5.546 | 523419 | 0.3388 | 0.3388 | 112.9 | 5.542 | 137780 |
| Medium_QC_2 | QC | 0.5 | 11/14/2023 16:08 | 0.5 | 5.535 | 746629 | 0.4809 | 0.4809 | 96.2 | 5.535 | 140196 |
| High_QC_2 | QC | 0.8 | 11/14/2023 16:19 | 0.8 | 5.539 | 1247792 | 0.8206 | 0.8206 | 102.6 | 5.539 | 138993 |
| preblank7 | DoubleBlank | | 11/14/2023 16:30 | | 5.545 | 304821 | 39.0905 | 39.0905 | | 5.603 | 725 |
| preblank118 | DoubleBlank | | 11/14/2023 16:40 | | 5.543 | 329077 | 10.771 | 10.771 | | 5.538 | 2838 |
| LLOQ_3 | QC | 0.05 | 11/14/2023 16:51 | 0.05 | 5.543 | 96861 | 0.0549 | 0.0549 | 109.9 | 5.543 | 129639 |
| Low_QC_3 | QC | 0.3 | 11/14/2023 17:01 | 0.3 | 5.541 | 537027 | 0.3735 | 0.3735 | 124.5 | 5.541 | 128726 |
| Medium_QC_3 | QC | 0.5 | 11/14/2023 17:12 | 0.5 | 5.542 | 762244 | 0.5538 | 0.5538 | 110.8 | 5.542 | 124768 |
| High_QC_3 | QC | 0.8 | 11/14/2023 17:23 | 0.8 | 5.542 | 1173848 | 0.8038 | 0.8038 | 100.5 | 5.542 | 133445 |
| preblank9 | DoubleBlank | | 11/14/2023 17:33 | | 5.545 | 315277 | 7.3987 | 7.3987 | | 5.532 | 3956 |
| preblank10 | DoubleBlank | | 11/14/2023 17:44 | | | | | | | 5.744 | 146 |
| LLOQ_4 | QC | 0.05 | 11/14/2023 17:55 | 0.05 | 5.546 | 97232 | 0.0522 | 0.0522 | 104.4 | 5.542 | 135469 |
| Low_QC_4 | QC | 0.3 | 11/14/2023 18:05 | 0.3 | 5.541 | 497136 | 0.3183 | 0.3183 | 106.1 | 5.541 | 138948 |
| Medium_QC_4 | QC | 0.5 | 11/14/2023 18:16 | 0.5 | 5.543 | 772421 | 0.5217 | 0.5217 | 104.3 | 5.543 | 133988 |
| High_QC_4 | QC | 0.8 | 11/14/2023 18:27 | 0.8 | 5.543 | 1175379 | 0.8344 | 0.8344 | 104.3 | 5.543 | 128794 |
| preblank11 | DoubleBlank | | 11/14/2023 18:37 | | 5.546 | 319913 | 11.3496 | 11.3496 | | 5.601 | 2619 |
| preblank12 | DoubleBlank | | 11/14/2023 18:48 | | 5.549 | 249653 | 151.5648 | 151.5648 | | 5.729 | 153 |
| LLOQ_5 | QC | 0.05 | 11/14/2023 18:59 | 0.05 | 5.543 | 89380 | 0.0508 | 0.0508 | 101.6 | 5.543 | 127233 |
| Low_QC_5 | QC | 0.3 | 11/14/2023 19:09 | 0.3 | 5.545 | 485334 | 0.3334 | 0.3334 | 111.1 | 5.545 | 129737 |
| Medium_QC_5 | QC | 0.5 | 11/14/2023 19:20 | 0.5 | 5.544 | 760813 | 0.4944 | 0.4944 | 98.9 | 5.544 | 139062 |
| High_QC_5 | QC | 0.8 | 11/14/2023 19:31 | 0.8 | 5.542 | 1143868 | 0.7666 | 0.7666 | 95.8 | 5.542 | 136219 |
| preblank13 | DoubleBlank | | 11/14/2023 19:41 | | 5.544 | 325806 | 21.746 | 21.746 | | 5.607 | 1393 |
| preblank14 | DoubleBlank | | 11/14/2023 19:52 | | 5.543 | 323687 | | | | | |
| LLOQ_6 | QC | 0.05 | 11/14/2023 20:03 | 0.05 | 5.545 | 86719 | 0.0446 | 0.0446 | 89.3 | 5.545 | 136226 |
| Low_QC_6 | QC | 0.3 | 11/14/2023 20:13 | 0.3 | 5.543 | 504615 | 0.3068 | 0.3068 | 102.3 | 5.543 | 146088 |
| Medium_QC_6 | QC | 0.5 | 11/14/2023 20:24 | 0.5 | 5.543 | 743547 | 0.5051 | 0.5051 | 101 | 5.543 | 133111 |
| High_QC_6 | QC | 0.8 | 11/14/2023 20:34 | 0.8 | 5.546 | 1238645 | 0.779 | 0.779 | 97.4 | 5.546 | 145204 |
| preblank15 | DoubleBlank | | 11/14/2023 20:45 | | 5.545 | 333317 | 29.7634 | 29.7634 | | 5.566 | 1041 |
| preblank16 | DoubleBlank | | 11/14/2023 20:56 | | 5.543 | 323483 | 10.0526 | 10.0526 | | 5.568 | 2989 |
| stab_dev_L1_-80C | Sample | 0.3 | 11/14/2023 21:06 | 0.3 | 5.545 | 457329 | 0.2623 | 0.2623 | 87.4 | 5.54 | 153653 |
| stab_dev_H1_-80C | Sample | 0.8 | 11/14/2023 21:17 | 0.8 | 5.545 | 964300 | 0.5349 | 0.5349 | 66.9 | 5.541 | 163263 |
| preblank17 | DoubleBlank | | 11/14/2023 21:28 | | | | | | | 5.751 | 193 |
| preblank18 | DoubleBlank | | 11/14/2023 21:39 | | 5.547 | 281317 | 32.2046 | 32.2046 | | 5.375 | 812 |
| stab_dev_L1_-20C | Sample | 0.3 | 11/14/2023 21:50 | 0.3 | 5.542 | 227913 | 0.1241 | 0.1241 | 41.4 | 5.542 | 152927 |
| stab_dev_H1_-20C | Sample | 0.8 | 11/14/2023 22:00 | 0.8 | 5.543 | 731297 | 0.404 | 0.404 | 50.5 | 5.543 | 162522 |
| preblank19 | DoubleBlank | | 11/14/2023 22:11 | | 5.545 | 332674 | 60.7118 | 60.7118 | | 5.532 | 510 |
| preblank20 | DoubleBlank | | 11/15/2023 8:07 | | 5.544 | 233276 | 15.5693 | 15.5693 | | 5.636 | 1393 |
| stab_dev_L1_4C | Sample | 0.3 | 11/15/2023 8:18 | 0.3 | 5.542 | 335406 | 0.2048 | 0.2048 | 68.3 | 5.542 | 142217 |
| stab_dev_H1_4C | Sample | 0.8 | 11/15/2023 8:29 | 0.8 | 5.545 | 1032043 | 0.6216 | 0.6216 | 77.7 | 5.541 | 150920 |
| preblank21 | DoubleBlank | | 11/15/2023 8:40 | | 5.545 | 320490 | 80.1748 | 80.1748 | | 5.566 | 372 |

| | | | | hydrazine Method | hydrazine Results | | | | | $^{15}N_2$-hydrazine (ISTD) Results | |
|---|---|---|---|---|---|---|---|---|---|---|---|
| Name | Type | Level | Acq. Date-Time | Exp. Conc. | RT | Resp. | Calc. Conc. | Final Conc. | Accuracy | RT | Resp. |
| preblank22 | DoubleBlank | | 11/15/2023 8:50 | | 5.544 | 315588 | 10.8319 | 10.8319 | | 5.637 | 2707 |
| stab_dev_L1_auto | Sample | 0.3 | 11/15/2023 9:01 | 0.3 | 5.546 | 448213 | 0.2689 | 0.2689 | 89.6 | 5.541 | 147112 |
| stab_dev_H1_auto | Sample | 0.8 | 11/15/2023 9:12 | 0.8 | 5.544 | 1152557 | 0.6748 | 0.6748 | 84.3 | 5.544 | 155539 |
| preblank23 | DoubleBlank | | 11/15/2023 9:23 | | 5.543 | 304844 | 3.03 | 3.03 | | 5.681 | 9315 |
| preblank24 | DoubleBlank | | 11/15/2023 9:34 | | 5.544 | 317121 | 33.3025 | 33.3025 | | 5.645 | 885 |
| stab_dev_L1_RT | Sample | 0.3 | 11/15/2023 9:44 | 0.3 | 5.546 | 394550 | 0.2419 | 0.2419 | 80.6 | 5.542 | 143125 |
| stab_dev_H1_RT | Sample | 0.8 | 11/15/2023 9:55 | 0.8 | 5.546 | 1023126 | 0.6052 | 0.6052 | 75.6 | 5.546 | 153586 |
| preblank25 | DoubleBlank | | 11/15/2023 10:06 | | 5.544 | 333072 | | | | | |
| preblank26 | DoubleBlank | | 11/15/2023 10:17 | | 5.545 | 327790 | | | | | |
| stab_dev_L2_-80C | Sample | 0.3 | 11/15/2023 10:27 | 0.3 | 5.546 | 462574 | 0.2735 | 0.2735 | 91.2 | 5.541 | 149373 |
| stab_dev_H2_-80C | Sample | 0.8 | 11/15/2023 10:38 | 0.8 | 5.542 | 1157913 | 0.7237 | 0.7237 | 90.5 | 5.542 | 145903 |
| preblank27 | DoubleBlank | | 11/15/2023 10:49 | | 5.545 | 312333 | 21.4092 | 21.4092 | | 5.659 | 1356 |
| preblank28 | DoubleBlank | | 11/15/2023 11:00 | | 5.545 | 327679 | 169.9098 | 169.9098 | | 5.512 | 179 |
| stab_dev_L2_-20C | Sample | 0.3 | 11/15/2023 11:11 | 0.3 | 5.543 | 300944 | 0.1717 | 0.1717 | 57.2 | 5.543 | 150300 |
| stab_dev_H2_-20C | Sample | 0.8 | 11/15/2023 11:21 | 0.8 | 5.545 | 695705 | 0.4406 | 0.4406 | 55.1 | 5.545 | 142182 |
| preblank29 | DoubleBlank | | 11/15/2023 11:32 | | 5.548 | 285698 | 109.0292 | 109.0292 | | 5.38 | 244 |
| preblank30 | DoubleBlank | | 11/15/2023 11:43 | | 5.543 | 328135 | 8.2133 | 8.2133 | | 5.535 | 3710 |
| stab_dev_L2_4C | Sample | 0.3 | 11/15/2023 11:54 | 0.3 | 5.545 | 355494 | 0.1839 | 0.1839 | 61.3 | 5.541 | 166664 |
| stab_dev_H2_4C | Sample | 0.8 | 11/15/2023 12:05 | 0.8 | 5.545 | 988840 | 0.5324 | 0.5324 | 66.5 | 5.545 | 168189 |
| preblank31 | DoubleBlank | | 11/15/2023 12:15 | | 5.542 | 328461 | | | | | |
| preblank32 | DoubleBlank | | 11/15/2023 12:26 | | 5.544 | 327960 | 34.2677 | 34.2677 | | 5.603 | 890 |
| stab_dev_L2_auto | Sample | 0.3 | 11/15/2023 12:37 | 0.3 | 5.544 | 467238 | 0.2652 | 0.2652 | 88.4 | 5.544 | 155361 |
| stab_dev_H2_auto | Sample | 0.8 | 11/15/2023 12:48 | 0.8 | 5.545 | 1163646 | 0.6757 | 0.6757 | 84.5 | 5.541 | 156837 |
| preblank33 | DoubleBlank | | 11/15/2023 12:58 | | 5.544 | 327129 | 15.5763 | 15.5763 | | 5.544 | 1952 |
| preblank34 | DoubleBlank | | 11/15/2023 13:09 | | 5.542 | 318176 | 66.7781 | 66.7781 | | 5.609 | 443 |
| stab_dev_L2_RT | Sample | 0.3 | 11/15/2023 13:20 | 0.3 | 5.546 | 413744 | 0.2564 | 0.2564 | 85.5 | 5.542 | 142036 |
| stab_dev_H2_RT | Sample | 0.8 | 11/15/2023 13:31 | 0.8 | 5.546 | 1011186 | 0.6598 | 0.6598 | 82.5 | 5.542 | 139501 |
| preblank35 | DoubleBlank | | 11/15/2023 13:42 | | 5.545 | 320574 | 25.9352 | 25.9352 | | 5.537 | 1149 |
| preblank36 | DoubleBlank | | 11/15/2023 13:53 | | | | | | | 5.731 | 363 |
| stab_dev_L3_-80C | Sample | 0.3 | 11/15/2023 14:03 | 0.3 | 5.545 | 398355 | 0.2254 | 0.2254 | 75.1 | 5.545 | 154405 |
| stab_dev_H3_-80C | Sample | 0.8 | 11/15/2023 14:14 | 0.8 | 5.543 | 1131600 | 0.6882 | 0.6882 | 86 | 5.543 | 149791 |
| preblank37 | DoubleBlank | | 11/15/2023 14:25 | | 5.552 | 287206 | 364.9618 | 364.9618 | | 5.384 | 73 |
| preblank38 | DoubleBlank | | 11/15/2023 14:36 | | 5.787 | 5883 | 0.9614 | 0.9614 | | 5.745 | 561 |
| stab_dev_L3_-20C | Sample | 0.3 | 11/15/2023 14:46 | 0.3 | 5.549 | 156806 | 0.0916 | 0.0916 | 30.5 | 5.544 | 137453 |
| stab_dev_H3_-20C | Sample | 0.8 | 11/15/2023 14:57 | 0.8 | 5.542 | 779429 | 0.4831 | 0.4831 | 60.4 | 5.542 | 145685 |
| preblank39 | DoubleBlank | | 11/15/2023 15:08 | | 5.545 | 318855 | | | | | |
| preblank40 | DoubleBlank | | 11/15/2023 15:19 | | 5.545 | 312974 | 163.579 | 163.579 | | 5.549 | 178 |
| stab_dev_L3_4C | Sample | 0.3 | 11/15/2023 15:29 | 0.3 | 5.545 | 393946 | 0.2411 | 0.2411 | 80.4 | 5.54 | 143327 |
| stab_dev_H3_4C | Sample | 0.8 | 11/15/2023 15:40 | 0.8 | 5.542 | 860110 | 0.4878 | 0.4878 | 61 | 5.542 | 159291 |
| preblank41 | DoubleBlank | | 11/15/2023 15:51 | | 5.545 | 314295 | | | | | |
| preblank42 | DoubleBlank | | 11/15/2023 16:02 | | 5.546 | 315599 | 10.7607 | 10.7607 | | 5.579 | 2725 |
| stab_dev_L3_auto | Sample | 0.3 | 11/15/2023 16:13 | 0.3 | 5.543 | 447607 | 0.2661 | 0.2661 | 88.7 | 5.543 | 148352 |
| stab_dev_H3_auto | Sample | 0.8 | 11/15/2023 16:23 | 0.8 | 5.545 | 1153001 | 0.6988 | 0.6988 | 87.4 | 5.541 | 150362 |
| preblank43 | DoubleBlank | | 11/15/2023 16:34 | | 5.546 | 239400 | 252.7856 | 252.7856 | | 5.731 | 88 |
| preblank44 | DoubleBlank | | 11/15/2023 16:45 | | 5.543 | 298807 | | | | | |
| stab_dev_L3_RT | Sample | 0.3 | 11/15/2023 16:56 | 0.3 | 5.545 | 401532 | 0.257 | 0.257 | 85.7 | 5.545 | 137551 |
| stab_dev_H3_RT | Sample | 0.8 | 11/15/2023 17:07 | 0.8 | 5.546 | 1037050 | 0.6744 | 0.6744 | 84.3 | 5.546 | 140027 |
| preblank45 | DoubleBlank | | 11/15/2023 17:17 | | 5.542 | 277469 | 22.8785 | 22.8785 | | 5.614 | 1128 |
| preblank46 | DoubleBlank | | 11/15/2023 17:28 | | 5.66 | 604 | 0.0672 | 0.0672 | | 5.656 | 687 |
| stab_stock_L1_-80C | Sample | 0.3 | 11/15/2023 17:39 | 0.3 | 5.547 | 467902 | 0.2879 | 0.2879 | 96 | 5.542 | 143900 |
| stab_stock_H1_-80C | Sample | 0.8 | 11/15/2023 17:50 | 0.8 | 5.543 | 1122218 | 0.7526 | 0.7526 | 94.1 | 5.543 | 136086 |
| preblank47 | DoubleBlank | | 11/15/2023 18:00 | | 5.653 | 987 | 0 | 0 | | 5.641 | 8228 |
| preblank48 | DoubleBlank | | 11/15/2023 18:11 | | 5.704 | 921 | 0.2031 | 0.2031 | | 5.532 | 394 |
| stab_stock_L1_-20C | Sample | 0.3 | 11/15/2023 18:22 | 0.3 | 5.542 | 192028 | 0.1144 | 0.1144 | 38.1 | 5.542 | 138523 |
| stab_stock_H1_-20C | Sample | 0.8 | 11/15/2023 18:33 | 0.8 | 5.545 | 1171363 | 0.7639 | 0.7639 | 95.5 | 5.545 | 139989 |
| preblank49 | DoubleBlank | | 11/15/2023 18:43 | | 5.527 | 10340 | 1.1718 | 1.1718 | | 5.548 | 811 |
| preblank50_ | DoubleBlank | | 11/16/2023 9:14 | | 5.704 | 518 | 0.0408 | 0.0408 | | 5.734 | 870 |
| stab_stock_L1_4C | Sample | 0.3 | 11/16/2023 9:25 | 0.3 | 5.544 | 370704 | 0.265 | 0.265 | 88.3 | 5.544 | 123376 |
| stab_stock_H1_4C | Sample | 0.8 | 11/16/2023 9:36 | 0.8 | 5.543 | 1007319 | 0.7934 | 0.7934 | 99.2 | 5.543 | 115979 |
| preblank51_ | DoubleBlank | | 11/16/2023 9:46 | | 5.595 | 653 | 0.0385 | 0.0385 | | 5.654 | 1143 |
| preblank52_ | DoubleBlank | | 11/16/2023 9:57 | | 5.556 | 573 | 0.0444 | 0.0444 | | 5.585 | 905 |
| stab_stock_L1_auto | Sample | 0.3 | 11/16/2023 10:08 | 0.3 | 5.541 | 481994 | 0.3182 | 0.3182 | 106.1 | 5.541 | 134740 |
| stab_stock_H1_auto | Sample | 0.8 | 11/16/2023 10:19 | 0.8 | 5.544 | 1248529 | 0.8893 | 0.8893 | 111.2 | 5.54 | 128500 |
| preblank53_ | DoubleBlank | | 11/16/2023 10:29 | | | | | | | 5.534 | 323 |
| preblank54_ | DoubleBlank | | 11/16/2023 10:40 | | | | | | | | |
| stab_stock_L1_RT | Sample | 0.3 | 11/16/2023 10:51 | 0.3 | 5.543 | 324980 | 0.2322 | 0.2322 | 77.4 | 5.543 | 122501 |
| stab_stock_H1_RT | Sample | 0.8 | 11/16/2023 11:02 | 0.8 | 5.541 | 1083639 | 0.7332 | 0.7332 | 91.6 | 5.541 | 134822 |
| preblank55_ | DoubleBlank | | 11/16/2023 11:13 | | 5.545 | 3132 | | | | | |
| preblank56_ | DoubleBlank | | 11/16/2023 11:24 | | 5.547 | 2023 | 0.0587 | 0.0587 | | 5.48 | 2568 |
| stab_stock_L2_-80C | Sample | 0.3 | 11/16/2023 11:34 | 0.3 | 5.546 | 387592 | 0.2762 | 0.2762 | 92.1 | 5.542 | 124007 |
| stab_stock_H2_-80C | Sample | 0.8 | 11/16/2023 11:45 | 0.8 | 5.542 | 955554 | 0.7617 | 0.7617 | 95.2 | 5.542 | 114518 |
| preblank57_ | DoubleBlank | | 11/16/2023 11:56 | | | | | | | 5.577 | 2349 |
| preblank58_ | DoubleBlank | | 11/16/2023 12:07 | | 5.554 | 1713 | 0.0731 | 0.0731 | | 5.596 | 1817 |
| stab_stock_L2_-20C | Sample | 0.3 | 11/16/2023 12:17 | 0.3 | 5.542 | 358733 | 0.2611 | 0.2611 | 87 | 5.542 | 121064 |
| stab_stock_H2_-20C | Sample | 0.8 | 11/16/2023 12:28 | 0.8 | 5.542 | 803697 | 0.6258 | 0.6258 | 78.2 | 5.542 | 116762 |
| preblank59_ | DoubleBlank | | 11/16/2023 12:39 | | 5.911 | 3759 | 0.0357 | 0.0357 | | 5.731 | 6961 |
| preblank60_ | DoubleBlank | | 11/16/2023 12:50 | | 5.55 | 968 | 0.022 | 0.022 | | 5.508 | 2460 |
| stab_stock_L2_4C | Sample | 0.3 | 11/16/2023 13:00 | 0.3 | 5.544 | 403506 | 0.3146 | 0.3146 | 104.9 | 5.544 | 114021 |
| stab_stock_H2_4C | Sample | 0.8 | 11/16/2023 13:11 | 0.8 | 5.541 | 823789 | 0.7787 | 0.7787 | 97.3 | 5.541 | 96607 |
| preblank61_ | DoubleBlank | | 11/16/2023 13:22 | | 5.553 | 2709 | | | | | |
| preblank62_ | DoubleBlank | | 11/16/2023 13:33 | | 5.544 | 1714 | 0.3228 | 0.3228 | | 5.565 | 473 |
| stab_stock_L2_auto | Sample | 0.3 | 11/16/2023 13:43 | 0.3 | 5.541 | 399632 | 0.318 | 0.318 | 106 | 5.541 | 111783 |
| stab_stock_H2_auto | Sample | 0.8 | 11/16/2023 13:54 | 0.8 | 5.545 | 828386 | 0.8848 | 0.8848 | 110.6 | 5.545 | 85688 |
| preblank63_ | DoubleBlank | | 11/16/2023 14:05 | | 5.555 | 2333 | 0.1853 | 0.1853 | | 5.521 | 1086 |
| preblank64_ | DoubleBlank | | 11/16/2023 14:16 | | 5.32 | 394 | 0.1406 | 0.1406 | | 5.379 | 236 |

| | | | | hydrazine Method | hydrazine Results | | | | | $^{15}N_2$-hydrazine (ISTD) Results | |
|---|---|---|---|---|---|---|---|---|---|---|---|
| **Name** | **Type** | **Level** | **Acq. Date-Time** | **Exp. Conc.** | **RT** | **Resp.** | **Calc. Conc.** | **Final Conc.** | **Accuracy** | **RT** | **Resp.** |
| stab_stock_L2_RT | Sample | 0.3 | 11/16/2023 14:27 | 0.3 | 5.542 | 328224 | 0.2258 | 0.2258 | 75.3 | 5.542 | 127024 |
| stab_stock_H2_RT | Sample | 0.8 | 11/16/2023 14:37 | 0.8 | 5.545 | 966537 | 0.7464 | 0.7464 | 93.3 | 5.541 | 118163 |
| preblank65_ | DoubleBlank | | 11/16/2023 14:48 | | 5.266 | 270 | 0.0434 | 0.0434 | | 5.38 | 433 |
| preblank66_ | DoubleBlank | | 11/16/2023 14:59 | | 5.707 | 302 | 0.0046 | 0.0046 | | 5.67 | 1463 |
| stab_stock_L3_-80C | Sample | 0.3 | 11/16/2023 15:10 | 0.3 | 5.543 | 403738 | 0.283 | 0.283 | 94.3 | 5.543 | 126235 |
| stab_stock_H3_-80C | Sample | 0.8 | 11/16/2023 15:21 | 0.8 | 5.544 | 749981 | 0.7871 | 0.7871 | 98.4 | 5.54 | 87029 |
| preblank67_ | DoubleBlank | | 11/16/2023 15:31 | | 5.532 | 2355 | | | | | |
| preblank78_ | DoubleBlank | | 11/16/2023 15:42 | | 5.537 | 4783 | 0.0914 | 0.0914 | | 5.562 | 4200 |
| stab_stock_L3_-20C | Sample | 0.3 | 11/16/2023 15:53 | 0.3 | 5.544 | 306979 | 0.2507 | 0.2507 | 83.6 | 5.54 | 107650 |
| stab_stock_H3_-20C | Sample | 0.8 | 11/16/2023 16:04 | 0.8 | 5.544 | 398756 | 0.5757 | 0.5757 | 72 | 5.54 | 62847 |
| preblank69_ | DoubleBlank | | 11/16/2023 16:14 | | 5.567 | 1808 | | | | | |
| preblank70_ | DoubleBlank | | 11/16/2023 16:25 | | 5.566 | 1422 | 1.4701 | 1.4701 | | 5.579 | 89 |
| stab_stock_L3_4C | Sample | 0.3 | 11/16/2023 16:36 | 0.3 | 5.543 | 396910 | 0.2806 | 0.2806 | 93.5 | 5.543 | 125114 |
| stab_stock_H3_4C | Sample | 0.8 | 11/16/2023 16:47 | 0.8 | 5.543 | 948181 | 0.8589 | 0.8589 | 107.4 | 5.543 | 100990 |
| preblank71_ | DoubleBlank | | 11/16/2023 16:58 | | 5.541 | 3033 | 0.2382 | 0.2382 | | 5.634 | 1116 |
| preblank72_ | DoubleBlank | | 11/16/2023 17:08 | | 5.546 | 5792 | | | | | |
| stab_stock_L3_auto | Sample | 0.3 | 11/16/2023 17:19 | 0.3 | 5.542 | 428179 | 0.2852 | 0.2852 | 95.1 | 5.542 | 132876 |
| stab_stock_H3_auto | Sample | 0.8 | 11/16/2023 17:30 | 0.8 | 5.542 | 1211829 | 0.8763 | 0.8763 | 109.5 | 5.542 | 126554 |
| preblank73_ | DoubleBlank | | 11/16/2023 17:41 | | 5.548 | 1727 | | | | | |
| preblank74_ | DoubleBlank | | 11/16/2023 17:52 | | 5.548 | 1810 | 0.0224 | 0.0224 | | 5.556 | 4549 |
| stab_stock_L3_RT | Sample | 0.3 | 11/16/2023 18:02 | 0.3 | 5.542 | 216915 | 0.1889 | 0.1889 | 63 | 5.542 | 99162 |
| stab_stock_H3_RT | Sample | 0.8 | 11/16/2023 18:13 | 0.8 | 5.546 | 1048267 | 0.7457 | 0.7457 | 93.2 | 5.541 | 128265 |
| preblank75_ | DoubleBlank | | 11/16/2023 18:24 | | 5.54 | 2403 | 1.7795 | 1.7795 | | 5.578 | 125 |
| preblank76_ | DoubleBlank | | 11/16/2023 18:35 | | 5.791 | 4792 | 0.2786 | 0.2786 | | 5.681 | 1521 |
| zerocalibrator2 | Blank | | 11/16/2023 18:45 | | 5.541 | 1401 | 0 | 0 | | 5.541 | 130235 |
| L1_2 | Cal | 0.05 | 11/16/2023 18:56 | 0.05 | 5.547 | 25877 | 0.0359 | 0.0359 | 71.8 | 5.543 | 47699 |
| L2_2 | Cal | 0.08 | 11/16/2023 19:07 | 0.08 | 5.543 | 81793 | 0.0786 | 0.0786 | 98.3 | 5.543 | 81649 |
| L3_2 | Cal | 0.1 | 11/16/2023 19:17 | 0.1 | 5.543 | 131546 | 0.1006 | 0.1006 | 100.6 | 5.543 | 106246 |
| L4_2 | Cal | 0.15 | 11/16/2023 19:28 | 0.15 | 5.544 | 379259 | 0.2818 | 0.2818 | 187.9 | 5.54 | 119029 |
| L5_2 | Cal | 0.2 | 11/16/2023 19:39 | 0.2 | 5.542 | 266372 | 0.1711 | 0.1711 | 85.6 | 5.542 | 133441 |
| L6_2 | Cal | 0.3 | 11/16/2023 19:49 | 0.3 | 5.544 | 135618 | 0.1339 | 0.1339 | 44.6 | 5.544 | 84979 |
| L7_2 | Cal | 0.4 | 11/16/2023 20:00 | 0.4 | 5.541 | 562164 | 0.4285 | 0.4285 | 107.1 | 5.541 | 118035 |
| L8_2 | Cal | 0.5 | 11/16/2023 20:10 | 0.5 | 5.545 | 676489 | 0.4933 | 0.4933 | 98.7 | 5.541 | 123905 |
| L9_2 | Cal | 0.7 | 11/16/2023 20:21 | 0.7 | 5.54 | 783049 | 0.6762 | 0.6762 | 96.6 | 5.54 | 105464 |
| L10_2 | Cal | 0.8 | 11/16/2023 20:32 | 0.8 | 5.545 | 1071116 | 0.7308 | 0.7308 | 91.4 | 5.545 | 133679 |
| L11_2 | Cal | 0.9 | 11/16/2023 20:42 | 0.9 | 5.543 | 713157 | 0.87 | 0.87 | 96.7 | 5.543 | 75005 |
| L12_2 | Cal | 1 | 11/16/2023 20:53 | 1 | 5.545 | 1326837 | 1.0363 | 1.0363 | 103.6 | 5.541 | 117459 |
| preblank77_ | DoubleBlank | | 11/16/2023 21:04 | | 5.549 | 6113 | 1.8363 | 1.8363 | | 5.457 | 307 |
| preblank78_ | DoubleBlank | | 11/16/2023 21:14 | | 5.302 | 256 | 0.3195 | 0.3195 | | 5.378 | 71 |
| **derivatization in various acids** | | | | | | | | | | | |
| blank_AA_1 | DoubleBlank | | 5/15/2024 16:52 | | 5.636 | 17310 | 0.6264 | 0.6264 | | 5.67 | 2681 |
| blank_AA_2 | DoubleBlank | | 5/15/2024 17:03 | | | | | | | 5.605 | 276 |
| blank_AA_3 | DoubleBlank | | 5/15/2024 17:14 | | 5.521 | 1089 | 0.1488 | 0.1488 | | 5.538 | 661 |
| zerocalbrator | Blank | | 5/15/2024 17:24 | | 5.534 | 3204 | 0 | 0 | | 5.534 | 70455 |
| L1 | Cal | 0.05 | 5/15/2024 17:35 | 0.05 | 5.539 | 44061 | 0.0498 | 0.0498 | 99.6 | 5.535 | 67833 |
| L2 | Cal | 0.08 | 5/15/2024 17:46 | 0.08 | 5.534 | 64672 | 0.0773 | 0.0773 | 96.6 | 5.53 | 69829 |
| L3 | Cal | 0.1 | 5/15/2024 17:56 | 0.1 | 5.533 | 74414 | 0.0876 | 0.0876 | 87.6 | 5.533 | 72277 |
| L4 | Cal | 0.15 | 5/15/2024 18:07 | 0.15 | 5.533 | 112711 | 0.147 | 0.147 | 98 | 5.538 | 69209 |
| L5 | Cal | 0.2 | 5/15/2024 18:18 | 0.2 | 5.537 | 152271 | 0.2022 | 0.2022 | 101.1 | 5.537 | 69729 |
| L6 | Cal | 0.3 | 5/15/2024 18:29 | 0.3 | 5.537 | 221429 | 0.3047 | 0.3047 | 101.6 | 5.537 | 68859 |
| L7 | Cal | 0.4 | 5/15/2024 18:39 | 0.4 | 5.543 | 187129 | 0.3799 | 0.3799 | 95 | 5.539 | 47090 |
| L8 | Cal | 0.5 | 5/15/2024 18:50 | 0.5 | 5.536 | 369964 | 0.525 | 0.525 | 105 | 5.536 | 68080 |
| L9 | Cal | 0.7 | 5/15/2024 19:01 | 0.7 | 5.536 | 517548 | 0.7147 | 0.7147 | 102.1 | 5.536 | 70460 |
| L10 | Cal | 0.8 | 5/15/2024 19:11 | 0.8 | 5.535 | 591378 | 0.854 | 0.854 | 106.7 | 5.535 | 67608 |
| L11 | Cal | 0.9 | 5/15/2024 19:22 | 0.9 | 5.536 | 657120 | 0.8575 | 0.8575 | 95.3 | 5.532 | 74823 |
| L12 | Cal | 1 | 5/15/2024 19:33 | 1 | 5.535 | 722340 | 0.9804 | 0.9804 | 98 | 5.535 | 72088 |
| blank_AA_4 | DoubleBlank | | 5/15/2024 19:43 | | 5.554 | 6958 | 0.5678 | 0.5678 | | 5.541 | 1186 |
| blank_AA_5 | DoubleBlank | | 5/15/2024 19:54 | | 5.564 | 4644 | 0.2211 | 0.2211 | | 5.535 | 1956 |
| 03_AA_1 | Sample | | 5/15/2024 20:05 | | 5.537 | 202806 | 0.277 | 0.277 | | 5.533 | 69035 |
| 03_AA_2 | Sample | | 5/15/2024 20:15 | | 5.536 | 207784 | 0.2872 | 0.2872 | | 5.536 | 68341 |
| 03_AA_3 | Sample | | 5/15/2024 20:26 | | 5.536 | 196329 | 0.2814 | 0.2814 | | 5.532 | 65841 |
| blank_AA_6 | DoubleBlank | | 5/15/2024 20:37 | | 5.532 | 2892 | 0.3727 | 0.3727 | | 5.536 | 741 |
| blank_AA_7 | DoubleBlank | | 5/15/2024 20:47 | | 5.752 | 1777 | 0.4105 | 0.4105 | | 5.584 | 415 |
| 08_AA_1 | Sample | | 5/15/2024 20:58 | | 5.533 | 581631 | 0.8261 | 0.8261 | | 5.533 | 68698 |
| 08_AA_2 | Sample | | 5/15/2024 21:09 | | 5.534 | 576026 | 0.8275 | 0.8275 | | 5.534 | 67926 |
| 08_AA_3 | Sample | | 5/15/2024 21:19 | | 5.537 | 559617 | 0.8144 | 0.8144 | | 5.537 | 67035 |
| blank_TFA_1 | DoubleBlank | | 5/15/2024 21:30 | | 5.546 | 2692 | 0.2418 | 0.2418 | | 5.727 | 1042 |
| blank_TFA_2 | DoubleBlank | | 5/15/2024 21:41 | | 5.56 | 1718 | 1.1156 | 1.1156 | | 5.383 | 151 |
| 03_TFA_1 | Sample | | 5/15/2024 21:51 | | 5.534 | 199553 | 0.2911 | 0.2911 | | 5.534 | 64809 |
| 03_TFA_2 | Sample | | 5/15/2024 22:02 | | 5.533 | 185084 | 0.2651 | 0.2651 | | 5.537 | 65693 |
| 03_TFA_3 | Sample | | 5/15/2024 22:12 | | 5.532 | 214078 | 0.3074 | 0.3074 | | 5.532 | 66004 |
| blank_TFA_3 | DoubleBlank | | 5/15/2024 22:23 | | 5.525 | 7141 | 0.8501 | 0.8501 | | 5.542 | 820 |
| blank_TFA_4 | DoubleBlank | | 5/15/2024 22:34 | | 5.543 | 4155 | 0.2455 | 0.2455 | | 5.623 | 1585 |
| 08_TFA_1 | Sample | | 5/15/2024 22:44 | | 5.534 | 592931 | 0.8397 | 0.8397 | | 5.534 | 68915 |
| 08_TFA_2 | Sample | | 5/15/2024 22:55 | | 5.532 | 578268 | 0.8081 | 0.8081 | | 5.532 | 69793 |
| 08_TFA_3 | Sample | | 5/15/2024 23:06 | | 5.538 | 569813 | 0.8321 | 0.8321 | | 5.529 | 66823 |
| blank_FA_1 | DoubleBlank | | 5/15/2024 23:16 | | 5.591 | 721 | 0.0585 | 0.0585 | | 5.579 | 978 |
| blank_FA_2 | DoubleBlank | | 5/15/2024 23:27 | | 5.533 | 2158 | 1.191 | 1.191 | | 5.538 | 178 |
| 03_FA_1 | Sample | | 5/15/2024 23:38 | | 5.534 | 212505 | 0.3073 | 0.3073 | | 5.53 | 65534 |
| 03_FA_2 | Sample | | 5/15/2024 23:48 | | 5.535 | 200466 | 0.2908 | 0.2908 | | 5.535 | 65159 |
| 03_FA_3 | Sample | | 5/15/2024 23:59 | | 5.535 | 201205 | 0.2541 | 0.2541 | | 5.531 | 74330 |
| blank_FA_3 | DoubleBlank | | 5/16/2024 0:10 | | 5.533 | 1554 | 0.4694 | 0.4694 | | 5.554 | 319 |
| blank_FA_4 | DoubleBlank | | 5/16/2024 0:20 | | 5.524 | 1425 | 0.2171 | 0.2171 | | 5.571 | 611 |

| | | | | hydrazine Method | hydrazine Results | | | | | $^{15}N_2$-hydrazine (ISTD) Results | |
|---|---|---|---|---|---|---|---|---|---|---|---|
| **Name** | **Type** | **Level** | **Acq. Date-Time** | **Exp. Conc.** | **RT** | **Resp.** | **Calc. Conc.** | **Final Conc.** | **Accuracy** | **RT** | **Resp.** |
| 08_FA_1 | Sample | | 5/16/2024 0:31 | | 5.536 | 616112 | 0.8691 | 0.8691 | | 5.532 | 69229 |
| 08_FA_2 | Sample | | 5/16/2024 0:42 | | 5.534 | 584745 | 0.7635 | 0.7635 | | 5.534 | 74618 |
| 08_FA_3 | Sample | | 5/16/2024 0:52 | | 5.535 | 568593 | 0.8102 | 0.8102 | | 5.53 | 68454 |
| blank1 | DoubleBlank | | 5/16/2024 1:03 | | 5.753 | 657 | 0.0138 | 0.0138 | | 5.56 | 2287 |
| blank2 | DoubleBlank | | 5/16/2024 1:14 | | 5.548 | 2577 | 0.1037 | 0.1037 | | 5.573 | 2162 |
| 03_1 | Sample | | 5/16/2024 1:24 | | 5.535 | 224601 | 0.2988 | 0.2988 | | 5.535 | 71149 |
| 03_2 | Sample | | 5/16/2024 1:35 | | 5.534 | 241075 | 0.3404 | 0.3404 | | 5.53 | 67423 |
| 03_3 | Sample | | 5/16/2024 1:46 | | 5.532 | 232180 | 0.3086 | 0.3086 | | 5.532 | 71321 |
| blank3 | DoubleBlank | | 5/16/2024 1:56 | | | | | | | 5.5 | 1777 |
| blank4 | DoubleBlank | | 5/16/2024 2:07 | | 5.642 | 1380 | 0.057 | 0.057 | | 5.629 | 1910 |
| 08_1 | Sample | | 5/16/2024 2:18 | | 5.532 | 613822 | 0.9099 | 0.9099 | | 5.532 | 65932 |
| 08_2 | Sample | | 5/16/2024 2:28 | | 5.535 | 626091 | 0.852 | 0.852 | | 5.531 | 71742 |
| 08_3 | Sample | | 5/16/2024 2:39 | | 5.535 | 626441 | 0.8792 | 0.8792 | | 5.535 | 69594 |
| blank5 | DoubleBlank | | 5/16/2024 2:49 | | 5.54 | 7710 | 2.1151 | 2.1151 | | 5.545 | 359 |
| blank6 | DoubleBlank | | 5/16/2024 3:00 | | 5.537 | 2409 | 0.2603 | 0.2603 | | 5.553 | 870 |
| **activity assay** | | | | | | | | | | | |
| blank1 | DoubleBlank | | 1/19/2024 17:20 | | 5.376 | 257 | 0.0088 | 0.0088 | | 5.397 | 1004 |
| blank_1 | DoubleBlank | | 1/19/2024 17:31 | | 5.523 | 2338 | 0.0027 | 0.0027 | | 5.536 | 13331 |
| zerocalibrator | Blank | | 1/19/2024 17:41 | | 5.516 | 841 | 0 | 0 | | 5.533 | 92346 |
| L1 | Cal | 0.05 | 1/19/2024 17:52 | 0.05 | 5.534 | 66313 | 0.0484 | 0.0484 | 96.9 | 5.534 | 85402 |
| L2 | Cal | 0.1 | 1/19/2024 18:02 | 0.1 | 5.535 | 138885 | 0.1087 | 0.1087 | 108.7 | 5.535 | 88605 |
| L3 | Cal | 0.3 | 1/19/2024 18:13 | 0.3 | 5.536 | 338552 | 0.2667 | 0.2667 | 88.9 | 5.54 | 92923 |
| L4 | Cal | 0.5 | 1/19/2024 18:24 | 0.5 | 5.535 | 626082 | 0.5332 | 0.5332 | 106.6 | 5.535 | 87621 |
| L5 | Cal | 0.8 | 1/19/2024 18:34 | 0.8 | 5.535 | 967239 | 0.8086 | 0.8086 | 101.1 | 5.535 | 89863 |
| L6 | Cal | 1 | 1/19/2024 18:45 | 1 | 5.536 | 1207744 | 0.9846 | 0.9846 | 98.5 | 5.536 | 92366 |
| blank6 | DoubleBlank | | 1/19/2024 23:01 | | 5.549 | 5959 | 0.0153 | 0.0153 | | 5.536 | 17486 |
| blank7 | DoubleBlank | | 1/19/2024 23:11 | | 5.546 | 3051 | 0.0051 | 0.0051 | | 5.537 | 14751 |
| replicate1-0 | Sample | | 1/19/2024 23:22 | | 5.537 | 42090 | 0.0246 | 0.0246 | | 5.537 | 90765 |
| replicate1-1 | Sample | | 1/19/2024 23:33 | | 5.537 | 483596 | 0.4138 | 0.4138 | | 5.537 | 86720 |
| replicate1-2 | Sample | | 1/19/2024 23:43 | | 5.54 | 654160 | 0.5664 | 0.5664 | | 5.544 | 86280 |
| replicate1-3 | Sample | | 1/19/2024 23:54 | | 5.54 | 830759 | 0.6655 | 0.6655 | | 5.54 | 93527 |
| replicate1-4 | Sample | | 1/20/2024 0:05 | | 5.538 | 879102 | 0.7494 | 0.7494 | | 5.538 | 88042 |
| replicate1-5 | Sample | | 1/20/2024 0:15 | | 5.542 | 1155329 | 0.9763 | 0.9763 | | 5.538 | 89102 |
| replicate1-6 | Sample | | 1/20/2024 0:26 | | 5.538 | 1339891 | 0.9518 | 0.9518 | | 5.538 | 105967 |
| replicate1-7 | Sample | | 1/20/2024 0:36 | | 5.541 | 1312545 | 1.0716 | 1.0716 | | 5.537 | 92312 |
| blank8 | DoubleBlank | | 1/20/2024 0:47 | | 5.621 | 5403 | 0.0149 | 0.0149 | | 5.533 | 16088 |
| blank9 | DoubleBlank | | 1/20/2024 0:58 | | 5.477 | 1006 | 0 | 0 | | 5.54 | 12194 |
| blank_2 | DoubleBlank | | 1/20/2024 7:54 | | 5.541 | 12730 | 0.0453 | 0.0453 | | 5.55 | 17321 |
| blank_3 | DoubleBlank | | 1/20/2024 8:05 | | 5.55 | 3346 | 0.0051 | 0.0051 | | 5.546 | 16180 |
| replicate2-0 | Sample | | 1/20/2024 8:15 | | 5.542 | 36019 | 0.0274 | 0.0274 | | 5.542 | 72029 |
| replicate2-1 | Sample | | 1/20/2024 8:26 | | 5.54 | 403054 | 0.3677 | 0.3677 | | 5.54 | 81079 |
| replicate2-2 | Sample | | 1/20/2024 8:37 | | 5.538 | 501072 | 0.4379 | 0.4379 | | 5.538 | 85027 |
| replicate2-3 | Sample | | 1/20/2024 8:48 | | 5.54 | 559204 | 0.5188 | 0.5188 | | 5.536 | 80390 |
| replicate2-4 | Sample | | 1/20/2024 8:58 | | 5.541 | 640323 | 0.6607 | 0.6607 | | 5.537 | 72594 |
| replicate2-5 | Sample | | 1/20/2024 9:09 | | 5.538 | 763234 | 0.7804 | 0.7804 | | 5.538 | 73441 |
| replicate2-6 | Sample | | 1/20/2024 9:20 | | 5.539 | 1128110 | 0.8346 | 0.8346 | | 5.539 | 101593 |
| replicate2-7 | Sample | | 1/20/2024 9:30 | | 5.54 | 869883 | 0.9307 | 0.9307 | | 5.54 | 70339 |
| blank_4 | DoubleBlank | | 1/20/2024 9:41 | | 5.54 | 3119 | 0.0006 | 0.0006 | | 5.544 | 21078 |
| blank_5 | DoubleBlank | | 1/20/2024 9:52 | | 5.545 | 3959 | 0.0058 | 0.0058 | | 5.537 | 18339 |
| blank_12 | DoubleBlank | | 1/20/2024 16:49 | | 5.54 | 8299 | 0.0235 | 0.0235 | | 5.536 | 18491 |
| blank_13 | DoubleBlank | | 1/20/2024 17:00 | | 5.548 | 8312 | 0.0289 | 0.0289 | | 5.535 | 15988 |
| replicate3-0 | Sample | | 1/20/2024 17:11 | | 5.542 | 16819 | 0.011 | 0.011 | | 5.537 | 59241 |
| replicate3-1 | Sample | | 1/20/2024 17:22 | | 5.54 | 250590 | 0.4243 | 0.4243 | | 5.54 | 43849 |
| replicate3-2 | Sample | | 1/20/2024 17:33 | | 5.541 | 468571 | 0.5069 | 0.5069 | | 5.541 | 68913 |
| replicate3-3 | Sample | | 1/20/2024 17:43 | | 5.541 | 445317 | 0.6515 | 0.6515 | | 5.537 | 51188 |
| replicate3-4 | Sample | | 1/20/2024 17:54 | | 5.54 | 665527 | 0.7088 | 0.7088 | | 5.536 | 70410 |
| replicate3-5 | Sample | | 1/20/2024 18:05 | | 5.542 | 764987 | 0.9784 | 0.9784 | | 5.542 | 58875 |
| replicate3-6 | Sample | | 1/20/2024 18:15 | | 5.543 | 489883 | 0.965 | 0.965 | | 5.543 | 38217 |
| replicate3-7 | Sample | | 1/20/2024 18:26 | | 5.541 | 1294811 | 1.0411 | 1.0411 | | 5.541 | 93709 |
| blank_14 | DoubleBlank | | 1/20/2024 18:37 | | 5.542 | 7312 | 0.0258 | 0.0258 | | 5.542 | 15253 |
| blank_15 | DoubleBlank | | 1/20/2024 18:47 | | 5.552 | 5042 | 0.0118 | 0.0118 | | 5.539 | 17066 |
| **hydrazine in *K. stuttgartiensis* cell lysate** | | | | | | | | | | | |
| blank1 | DoubleBlank | | 12/4/2023 14:59 | | 5.54 | 966 | 0.0588 | 0.0588 | | 5.616 | 1327 |
| blank2 | DoubleBlank | | 12/4/2023 15:10 | | 5.795 | 1666 | 0.0096 | 0.0096 | | 5.673 | 10175 |
| blank3 | DoubleBlank | | 12/4/2023 15:29 | | 5.606 | 1775 | | | | | |
| L1_1 | Cal | 0.05 | 12/4/2023 15:40 | 0.05 | 5.538 | 74709 | 0.046 | 0.046 | 92.1 | 5.538 | 128494 |
| L1_2 | Cal | 0.08 | 12/4/2023 15:50 | 0.08 | 5.537 | 126656 | 0.0776 | 0.0776 | 97 | 5.537 | 134245 |
| L1_3 | Cal | 0.1 | 12/4/2023 16:01 | 0.1 | 5.539 | 158420 | 0.0965 | 0.0965 | 96.5 | 5.535 | 136561 |
| L1_4 | Cal | 0.15 | 12/4/2023 16:12 | 0.15 | 5.536 | 256188 | 0.1694 | 0.1694 | 113 | 5.536 | 128326 |
| L1_5 | Cal | 0.2 | 12/4/2023 16:22 | 0.2 | 5.537 | 302166 | 0.1763 | 0.1763 | 88.1 | 5.537 | 145639 |
| L1_6 | Cal | 0.3 | 12/4/2023 16:33 | 0.3 | 5.537 | 464434 | 0.2969 | 0.2969 | 99 | 5.537 | 134298 |
| L1_7 | Cal | 0.4 | 12/4/2023 16:44 | 0.4 | 5.538 | 645216 | 0.4356 | 0.4356 | 108.9 | 5.538 | 127821 |
| L1_8 | Cal | 0.5 | 12/4/2023 16:54 | 0.5 | 5.537 | 727413 | 0.4886 | 0.4886 | 97.7 | 5.537 | 128618 |
| L1_9 | Cal | 0.7 | 12/4/2023 17:05 | 0.7 | 5.493 | 343298 | 0.6499 | 0.6499 | 92.8 | 5.497 | 45744 |
| L1_10 | Cal | 0.8 | 12/4/2023 17:16 | 0.8 | 5.536 | 806852 | 0.6885 | 0.6885 | 86.1 | 5.536 | 101520 |
| L1_11 | Cal | 0.9 | 12/4/2023 17:26 | 0.9 | 5.538 | 1206465 | 0.9126 | 0.9126 | 101.4 | 5.538 | 114715 |
| L1_12 | Cal | 1 | 12/4/2023 17:37 | 1 | 5.553 | 756699 | 1.0799 | 1.0799 | 108 | 5.549 | 60854 |
| blank4 | DoubleBlank | | 12/4/2023 17:48 | | 5.532 | 1655 | 0.3621 | 0.3621 | | 5.545 | 394 |
| blank5 | DoubleBlank | | 12/4/2023 17:58 | | 5.54 | 835 | 0.0427 | 0.0427 | | 5.632 | 1538 |
| blank6 | DoubleBlank | | 12/4/2023 18:41 | | 5.548 | 2698 | | | | | |
| blank7 | DoubleBlank | | 12/4/2023 18:52 | | | | | | | 5.603 | 1049 |

| | | | | hydrazine Method | hydrazine Results | | | | | $^{15}N_2$-hydrazine (ISTD) Results | |
|---|---|---|---|---|---|---|---|---|---|---|---|
| Name | Type | Level | Acq. Date-Time | Exp. Conc. | RT | Resp. | Calc. Conc. | Final Conc. | Accuracy | RT | Resp. |
| pellet_kuenenia_1 | Sample | | 12/4/2023 19:02 | | 5.542 | 293276 | 0.2569 | 0.2569 | | 5.538 | 97808 |
| pellet_kuenenia_2 | Sample | | 12/4/2023 19:13 | | 5.538 | 288042 | 0.2629 | 0.2629 | | 5.538 | 93901 |
| pellet_kuenenia_3 | Sample | | 12/4/2023 19:23 | | 5.538 | 285986 | 0.2318 | 0.2318 | | 5.542 | 105496 |
| blank8 | DoubleBlank | | 12/4/2023 19:34 | | 5.539 | 6628 | 2.9871 | 2.9871 | | 5.539 | 193 |
| blank9 | DoubleBlank | | 12/4/2023 19:45 | | 5.568 | 1542 | 0.034 | 0.034 | | 5.652 | 3478 |
| L2_1 | Cal | 0.05 | 12/4/2023 19:55 | 0.05 | 5.539 | 91922 | 0.0557 | 0.0557 | 111.4 | 5.539 | 132750 |
| L2_2 | Cal | 0.08 | 12/4/2023 20:06 | 0.08 | 5.541 | 116652 | 0.0732 | 0.0732 | 91.5 | 5.541 | 130693 |
| L2_3 | Cal | 0.1 | 12/4/2023 20:17 | 0.1 | 5.542 | 169962 | 0.1073 | 0.1073 | 107.3 | 5.542 | 132424 |
| L2_4 | Cal | 0.15 | 12/4/2023 20:27 | 0.15 | 5.542 | 235746 | 0.1489 | 0.1489 | 99.3 | 5.542 | 133863 |
| L2_5 | Cal | 0.2 | 12/4/2023 20:38 | 0.2 | 5.541 | 312265 | 0.203 | 0.203 | 101.5 | 5.541 | 131122 |
| L2_6 | Cal | 0.3 | 12/4/2023 20:49 | 0.3 | 5.54 | 435306 | 0.2892 | 0.2892 | 96.4 | 5.54 | 129213 |
| L2_7 | Cal | 0.4 | 12/4/2023 20:59 | 0.4 | 5.541 | 597146 | 0.4246 | 0.4246 | 106.1 | 5.541 | 121336 |
| L2_8 | Cal | 0.5 | 12/4/2023 21:10 | 0.5 | 5.54 | 773916 | 0.4901 | 0.4901 | 98 | 5.54 | 136425 |
| L2_9 | Cal | 0.7 | 12/4/2023 21:20 | 0.7 | 5.543 | 1166937 | 0.7433 | 0.7433 | 106.2 | 5.543 | 136076 |
| L2_10 | Cal | 0.8 | 12/4/2023 21:31 | 0.8 | 5.541 | 1210913 | 0.8156 | 0.8156 | 101.9 | 5.541 | 128760 |
| L2_11 | Cal | 0.9 | 12/4/2023 21:42 | 0.9 | 5.542 | 1385327 | 0.8712 | 0.8712 | 96.8 | 5.538 | 137947 |
| L2_12 | Cal | 1 | 12/4/2023 21:52 | 1 | 5.542 | 1492448 | 1.0297 | 1.0297 | 103 | 5.542 | 125845 |
| blank10 | DoubleBlank | | 12/4/2023 22:03 | | 5.782 | 351 | 0.0661 | 0.0661 | | 5.61 | 433 |
| blank11 | DoubleBlank | | 12/4/2023 22:14 | | | | | | | 5.735 | 2964 |
